# Pervasive *cis*-regulatory co-option of a transposable element family reinforces cell identity across the mouse immune system

**DOI:** 10.64898/2025.12.22.696042

**Authors:** Jason D. Chobirko, Cédric Feschotte, Andrew Grimson

## Abstract

Transposable elements (TEs) make up about half of the human and mouse genomes and play important regulatory roles in immune responses. However, the *cis*-regulatory contribution of TEs to immune cell development is less characterized. Here, we analyzed hundreds of chromatin accessibility, gene expression, transcription factor occupancy and DNA-DNA contact datasets in diverse mouse and human immune cells. We identified two rodent-specific TE subfamilies, ORR1E and ORR1D2 (collectively, ODE) that have transformed into cell type-specific enhancers across the mouse immune system. ODE loci acquired mutations post-insertion that enable differential binding of lineage-specifying transcription factors. ODEs show evidence of evolutionary sequence constraint and contact promoters of hundreds of genes in immune cells. ODE-targeted genes show cell type-specific and mouse-specific increases in expression compared to concordant human cell types. Thus, a single TE family can undergo functionalization after its genomic spread to generate batteries of cell-specific enhancers supporting a complex developmental cascade.

## Introduction

Transposable elements (TEs) are mobile elements capable of selfish propagation within genomes. TEs are found in nearly all organisms and make up a large percentage of the human (∼54%) [1] and mouse (∼40%) [2] genomes. To facilitate their propagation, TEs often harbor regulatory sequences – promoters and enhancers – with multiple transcription factor (TF) binding sites driving the expression of TE-encoded genes [3,4]. This reliance on host TFs for propagation has evolutionarily tied TEs to their hosts, leading to species-specific TE expansions and distinct repertoires across species [5]. Notably, a sizable fraction of TEs in human (∼24%) and mouse (∼32%) are unique to the primate and rodent lineages, respectively [2]. Thus, lineage-specific amplification of TEs make them compelling candidates to introduce *cis*-regulatory elements (CREs) that impact proximal gene expression, rewire networks and drive phenotypic changes between species [3,4,6]. However, most TE insertions have neutral or detrimental effects on host fitness [7–9]. Co-opted TEs often bring regulatory novelty to gene expression and their sequences exhibit signatures of evolutionary constraint, i.e., purifying selection [3,4,10–13]. TEs also drive enhancer turnover – species-specific shifting from an ancestral enhancer to a proximal novel enhancer [14–17]. However, what drives certain TE loci or families to get co-opted as CREs within biological processes or cell types during evolution remains poorly understood.

To investigate sequence determinants that facilitate co-option of TEs as CREs, we sought to identify TE families with signatures of co-option in a rapidly evolving process. The immune system is one such rapidly evolving system [18], consisting of diverse cell types, each playing distinct roles to protect against threats. Immune cells derive from progenitor hematopoietic stem cells (HSCs) through a differentiation cascade that has been extensively studied across multiple organisms [18–21]. Differences in immune gene regulatory networks between species underscore the rapid evolution of this system and the impact of regulatory novelty on immune function [22]. Previous studies have investigated the role of TEs across components of the immune system [23–35], implicating TEs as sources of regulatory novelty in the evolution and function of immune cells. However, the extent to which specific TEs have been co-opted as CREs across immune cell development itself is largely unknown [23].

We investigated the co-option of TEs as CREs across mouse immune cell development. We used a compendium of genomic datasets from mouse and human, including chromatin accessibility (DNase-seq and ATAC-seq), gene expression (RNA-seq), TF binding profiles (ChIP-seq), as well as DNA-DNA contact maps (Micro-C) that we generated from mouse CD8+ T cells. We discovered two rodent-specific TE subfamilies, <u>O</u>RR1<u>D</u>2 and ORR1<u>E</u> (henceforth collectively termed ODE), make a prominent contribution to CREs active throughout mouse immune cell development. Notably, these elements stratify by their cell type-specific accessibility patterns, unique combinations of cell type-specifying TF motifs and binding patterns for these factors. We implicate ODEs as “boosters” of gene expression in mouse, serving to increase cell-type specific expression of mouse genes relative to their human orthologs, and to incorporate mouse-specific genes into immune cell regulatory networks. ODEs with cell type-specific accessibility patterns had increased sequence conservation compared to other ODEs. Collectively, our data indicates that ODEs are a potent source of regulatory novelty in rodents, which appears to have primarily reinforced immune cell type identity.

## Results

### Subfamily-specific TE accessibility profiles reflect immune cell-type identities

To evaluate the *cis*-regulatory contribution of TEs throughout mouse immune cell development, we first reanalyzed 90 ATAC-seq data sets across 10 immune cell types [36], which include progenitors, differentiated and activated cells. Samples were grouped by cell identities: hematopoietic stem cells (HSCs, 5 samples), B cells (19), natural killer cells (NKs, 6), innate lymphoid cells (ILCs, 4), T cells (conventional αβ, 15; γδ, 7), activated T and NKT cells (Act T, 10), monocytes (Mo, 2), macrophages together with granulocytes (MFs, 12), dendritic cells (DCs, 5) and non-immune-derived stromal cells (5; Fig. 1A). No study has previously investigated the contribution of TEs to accessibility across this dataset. To quantify accessible TEs (as a proxy for *cis*-regulatory activity), we identified regions of increased accessibility over background (peaks) genome-wide across each sample. We intersected the summit of each peak against TE annotations, enriching for CREs derived from the TE rather than flanking sequence (see Methods). Of the ∼330,000 peaks identified, the majority (∼90%) are distal to promoters, or >2kb from an annotated transcription start site. We found that ∼3.2% (∼123,000) of TE loci are accessible in at least one sample, ranging from 2-9% when partitioned by TE class (Fig. S1A). The majority (∼117,000; ∼95%) of accessible TEs are distal to promoters, with 6.6-25.6% of peaks annotated as TE-derived, representing an underrepresentation compared to the genomic fraction of TEs (Fig. S1B; Table S1). Overall, the contribution of TEs to ATAC-seq peaks is comparable to previously estimated contributions to mammalian CREs in various contexts [15,37].

**Fig. 1.**
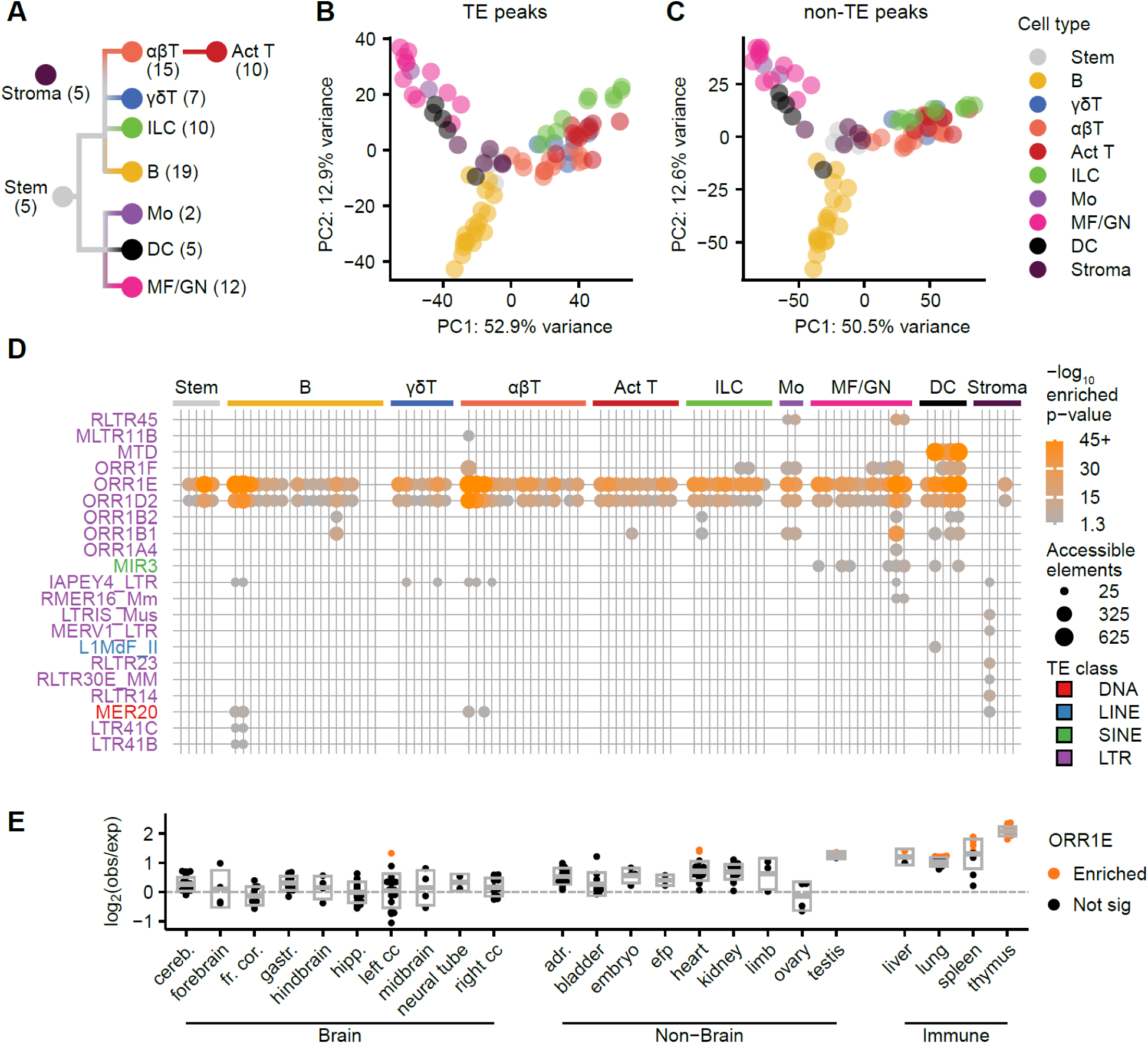
ODEs are enriched in accessibility across mouse immune cell development. (**A**) Cartoon depicting cell type annotation and number of ATAC-seq datasets across mouse immune cells. (**B – C**) PCAs of top 1,000 most variable TE-derived (B) or non-TE-derived (C) ATAC-seq unified peaks, with type identities color-coded. (**D**) Dotplot of enriched TE families significantly (permutation test<0.05 and Fisher’s Exact Test adjusted p-value<0.05) more accessible than shuffled peak background. (**E**) Enrichment of ORR1E subfamily DNase-seq peaks from Mouse Developmental Matrix split between tissue type, where each dot represents a dataset taken at different developmental timepoints. Orange dots indicate significant (permutation test<0.05 and Fisher’s Exact Test adjusted p-value<0.05) enrichment.

We sought to determine whether accessibility of TE-derived peaks alone could differentiate cell type-specific chromatin accessibility profiles across mouse immune cell development. We used the top 1,000 most-variable TE-derived (Fig. 1B) or non-TE-derived (Fig. 1C) peaks to perform principal component analysis. The results yield concordant partitioning of cells for TE-derived and non-TE-derived sets, with four clusters: 1) T cells, NKs, ILCs and activated adaptive cells; 2) monocytes, MFs and DCs; 3) B cells; and 4) HSCs and stromal cells. These clusters recapitulate expected similarities among these cell types [38]. Thus, accessible chromatin profiles at TEs alone resolve major immune cell lineages.

### ORR1D2 and ORR1E are accessible throughout immune cell development

Due to the distinct evolutionary origin, sequence and regulatory activities of different TEs [3,4,10], we hypothesized that certain subfamilies would exhibit distinctive chromatin accessibility patterns across immune cell development. Thus, we tested each subfamily for overrepresentation in accessibility outside of gene promoters across each ATAC-seq sample, controlling for subfamily copy number and genomic distribution. After filtering for significantly enriched TE subfamilies, we identified 21 subfamilies enriched in at least one sample (Fig. 1D). Most enriched subfamilies (∼86%) were long terminal repeats (LTRs) of endogenous retroviruses, which often contain *cis*-regulatory elements (CREs) [3,4]. Among enriched subfamilies, ORR1D2 and ORR1E stood out for their pervasive enrichment in accessibility across immune cell development, being enriched in 77 and 81 of 90 samples, respectively (Fig 1D).

ORR1D2 and ORR1E are older subfamilies of the rodent-specific origin-region repeat 1 (ORR1) family, and are the LTRs for elements of the mammalian apparent LTR transposon (MaLR) class [39,40]. Previously, ORR1E has been implicated as CREs in dendritic cells [41], and both ORR1D2 and ORR1E are enriched for accessible chromatin in CD8+ T cells [33] and monocytes [26]. However, the role of <u>O</u>RR1<u>D</u>2 and ORR1<u>E</u> (collectively, ODE) as CREs across immune cell development has not been investigated systematically. To determine whether the enrichment in accessibility of ODEs is unique to immune cell development or merely reflects non-specific accessibility, we analyzed 305 ATAC-seq and DNase-seq datasets across 27 tissues from the ENCODE mouse developmental matrix dataset [42,43]. Enrichment for ORR1E (Fig. 1E) and ORR1D2 (Fig. S1C-D) was maximal in tissues containing high concentrations of immune cells, such as the thymus and spleen, with low enrichment or even depletion in non-immune tissues such as brain. These findings highlight ODEs as candidate CREs both pervasive and specific to the immune system.

### ODE loci stratify by accessibility and TF motif enrichment

Typically, TE subfamilies act as CREs in distinct niches, a phenomenon mediated by unique combinations of TF binding sites across subfamilies [3,44–47]. We hypothesized that ODE loci either 1) exhibit broad accessibility across immune cells, or 2) contain subsets of elements that manifest with distinct, cell type-specific accessibility patterns. To test this, we calculated an accessibility score for all ORR1E (n=2,608) and ORR1D2 (n=1,346) loci that overlapped accessible peaks across any of the 90 ATAC-seq samples (Fig. 2A & S2A, respectively). Next, we performed unsupervised clustering of ODE loci by their accessibility scores and defined an optimal number of 8 clusters for ORR1E (Fig. S2B) and 6 for ORR1D2 (Fig. S2C), with 161 to 524 ODE loci per cluster partitioned by cell type-specific accessibility. Thus, ODE loci manifest distinct accessibility patterns across immune cells.

**Fig. 2.**
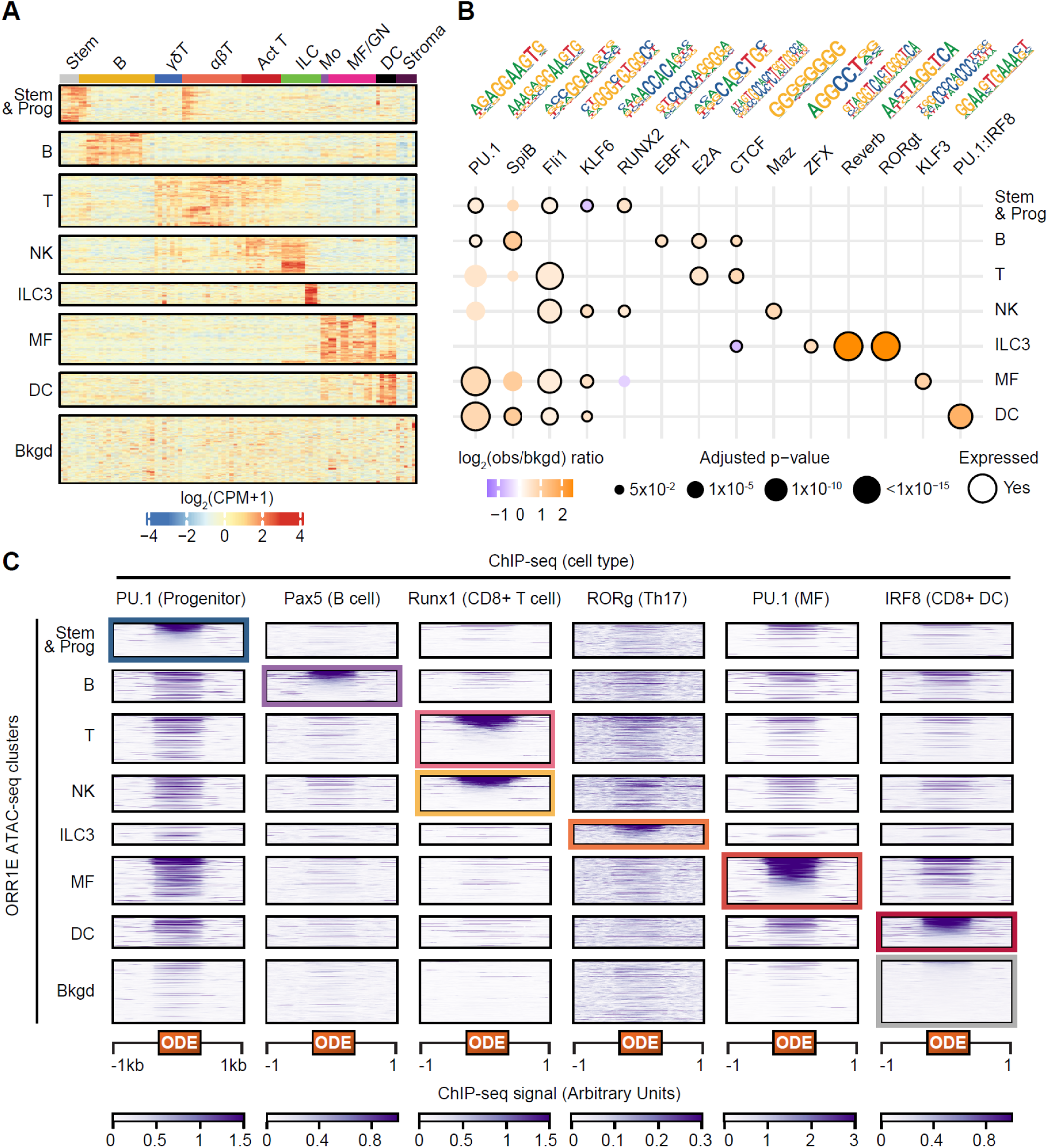
ODEs stratify by accessibility patterns, TF motif enrichment and TF binding. (**A**) Heatmap of 2,608 accessible ORR1E loci (rows) across ImmGen ATAC-seq data (columns) with kmeans groups split by rows; color indicates row-scaled quantile normalized log_2_(CPM + 1). (**B**) Significant (Binomial test adjusted p-value<0.05) motif enrichment results from HOMER on ORR1E clusters compared to the ‘background’ cluster from panel A. Dots sized to indicate adjusted p-value and outlined if the corresponding TF is expressed (>1 TPM) across the majority of cell types composing each cluster. Motif positional weight matrices are provided for each TF motif (**C**) Heatmaps of TF ChIP-seq datasets across ORR1E clusters expanded by 1kb upstream and downstream of individual ORR1E loci for each accessibility cluster. Highlighted boxes indicate dataset used for descending row sorting by signal intensity for each cluster across all ChIP-seq heatmaps.

To understand stratification of ODE loci across cell type-specific accessibility patterns, we examined the cell types corresponding to each ODE cluster. For example, cluster 1 contains ODE loci preferentially accessible in HSCs and progenitor T and B cells; thus we annotated this cluster “Stem & Prog” (Fig. 2A & S2A). Some clusters exhibit striking specificity for a given cell type, such as the “ILC3” cluster, which only contains ORR1E loci preferentially accessible in ILC3 cells. Similarly, one ORR1E cluster corresponds to macrophages (MF) and one cluster of ORR1D2 loci corresponds to macrophages and dendritic cells (MF & DC; which derive from a common progenitor [48]). For each subfamily, all but one cluster corresponds to distinct immune cell types and are depleted for accessibility in non-immune-derived stromal cells. The remaining cluster (cluster 6 for ORR1D2, 22.4% of loci; cluster 8 for ORR1E, 20.1% of loci) did not correspond to any obvious subset of cell types and therefore were labeled as “background” (Fig 2A & S2A).

We next sought to understand whether TF motifs correlated with the partitioning of ODEs across cell types by searching for overrepresented motifs per cluster relative to the “background” cluster of the same ODE subfamily. Out of 430 TF motifs searched, 59 (ORR1D2) and 93 (ORR1E) were enriched in at least one cluster. Using matched RNA-seq data [36], we filtered for TFs expressed in cell types matching cluster identity and visualized the top 3 enriched motifs for each ORR1E (Fig. 2B) and ORR1D2 (Fig. S2D) cluster. Enriched TF motifs recapitulated known key regulators of the cognate cell type identities. For example, the ETS motif was enriched in all clusters except for the “ILC3” cluster, an observation consistent with studies showing ETS family member PU.1 (Spi1) as necessary for HSC differentiation and establishing lineage-specific identities of multiple immune cells [49,50]. Conversely, the ILC3 cluster was enriched for ROR family motifs, such as RORgt, which are key TFs for ILC3 and Th17 development and for ILC3 effector function [51,52]. RORgt acts as both an activator of signature Th17 genes and a repressor of signature genes of other CD4+ T cell subsets [53]. The Stem & Prog and NK clusters were enriched for motifs of the Runx family, which are necessary for formation of HSCs and progenitor cells [54], cooperate with ETS-family members such as PU.1 in regulating gene expression [55], serve as master regulators of T cells [56], and facilitate robust activation of NK cells [57]. The B clusters were enriched for the EBF1 motif; indeed EBF1 is a master regulator of B cell development, necessary for survival and differentiation of progenitor B cells and maintenance of B cell identity [58–60]. In sum, ODE loci with cell type-specific accessibility patterns harbor distinct repertoires of cognate lineage-specifying TF motifs.

Our observation that ODEs are enriched for lineage-specifying TF motifs motivated us to ask whether ODEs are bound by these TFs in cell types concordant with their accessibility patterns. We analyzed ChIP-seq data for ODE-enriched TFs necessary for lineage-specific immune cell development, including PU.1 in progenitor HSCs [61] and bone marrow-derived macrophages [62], Pax5 in B cells [63], Runx1 in naïve CD8+ T cells [64], RORg in Th17s [65] and IRF8 in CD8+ DCs [66]. We overlayed ChIP-seq signal for accessible ORR1E (Fig. 2C) and ORR1D2 (Fig. S2E) loci and sorted each cluster by ChIP-seq signal. Each TF examined was preferentially bound to ODEs most accessible in the assayed cell type, such as IRF8 binding 51% of “DC” ORR1Es, and Pax5 binding ∼32% of “B” ORR1Es (Table S2). IRF8 is critical in controlling response to infection, maintenance and differentiation of dendritic cells from HSCs [67,68], and a combined motif of PU.1 and IRF8 was uniquely enriched in “DC” ORR1Es (Fig. 2B). Pax5 is necessary for establishing B cell identity and development [63]. “T” and “NK” ORR1Es had the strongest enrichment in RUNX1 binding (∼22% and ∼21%, respectively, Table S2), a key factor controlling T and NK cell development and function [57,69]. Similar TF binding preferences were observed for ORR1D2 clusters (Table S3 & Fig. S2E). These results suggest that ODE subsets are preferentially bound by lineage-specifying TFs across immune cell development concordant with their motif enrichments and accessibility patterns.

### ODEs are preferentially bound by pioneer factor PU.1 in progenitor cells

As ODEs are accessible across immune cell development, we hypothesized that they are bound by pioneer factors – TFs that initiate opening of chromatin [70,71]. Such pioneer factor binding could occur early in immune cell development, in HSCs, or later within differentiated cells. Of the TFs binding ODEs (Fig. 2B & S2D), PU.1 is conspicuous as a pioneer factor and master regulator throughout immune system development [72–74]. To determine whether PU.1 binding to ODEs in early progenitor cells was associated with binding of lineage-specifying TFs in differentiated cells, we annotated ODEs by their patterns of combined TF binding events across ORR1E (Fig. 3A) and ORR1D2 (Fig. S3A). All ODE clusters had a significantly higher percentage of elements bound by at least one TF compared to “background”, ranging between 31% and 75% for ORR1E (Fig. 3B) and between 46% and 70% for ORR1D2 (Fig. S3B). Except for the ORR1E “ILC3” cluster, all clusters have stronger propensity for being bound by at least two TFs compared to the “background” cluster (ORR1E, Fig. 3C; ORR1D2, Fig. S3C). Strikingly, the majority of ODE loci bound by at least two TFs are also bound by PU.1 in progenitor cells (∼90% for ORR1E, Fig. 3C; ∼83% for ORR1D2, Fig. S3C). These results implicate PU.1 binding to ODEs in progenitor cells as an early event promoting binding of lineage-specific TFs later in immune cell development.

**Fig. 3:**
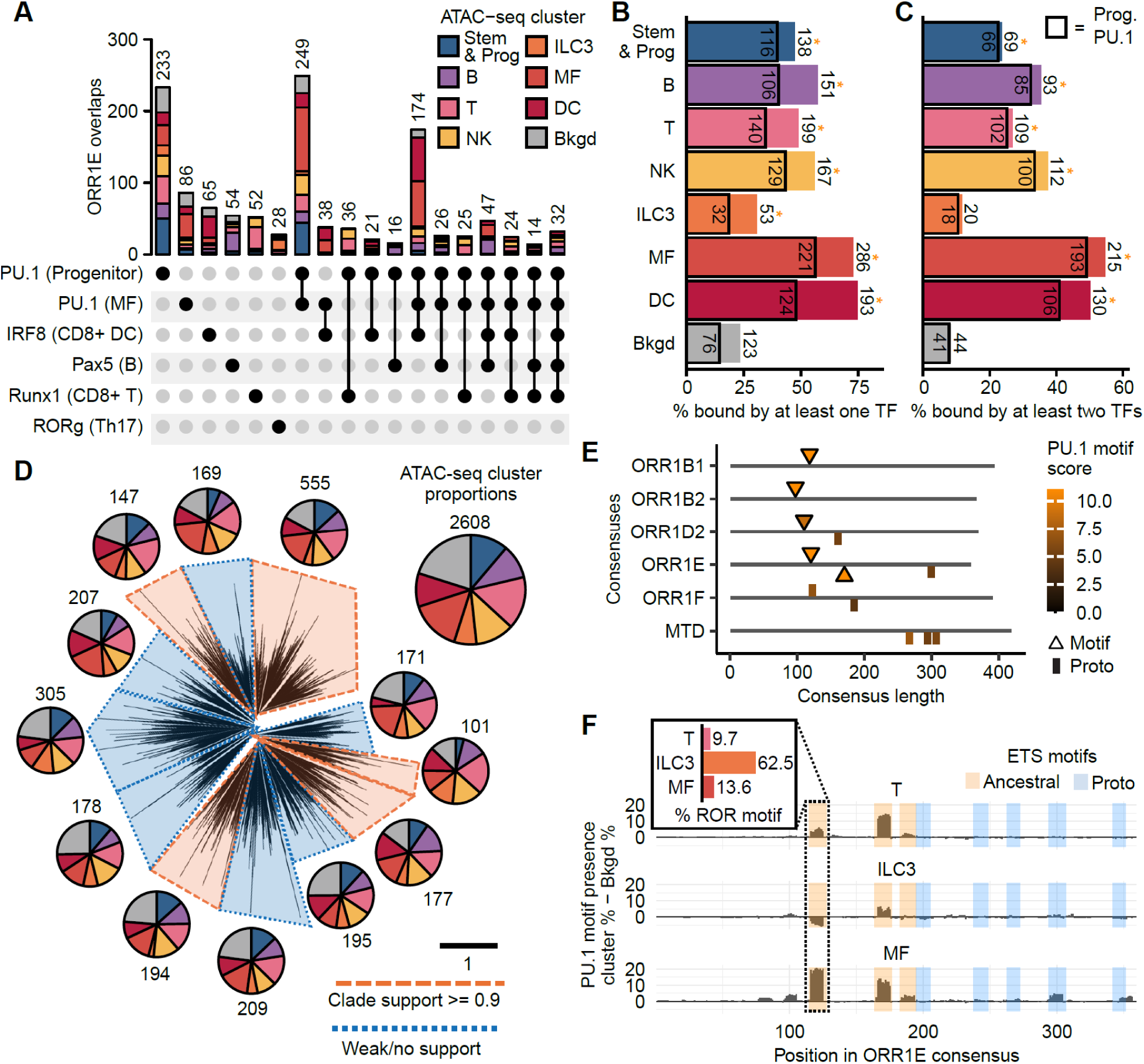
ODEs are preferentially bound by lineage-specifying TFs. (**A**) Upset plot showing proportions of ORR1E bound by any combination of TFs shown in Fig. 2C with at least 10 instances. (**B – C**) Barplots showing total proportion of loci per ORR1E cluster bound by at least one (B) or two (C) TFs in distinct cell types. Box indicates proportion bound by PU.1 in progenitor cells and * indicates significantly (Fisher’s Exact Test adjusted p-value<0.05) higher total % bound compared to “Bkgd” for non-boxed proportions. (**D**) Phylogenetic tree of accessible ORR1Es highlighting clade support (>= 0.90; in orange) or weak/no robust clade support (< 0.90); black scale bar denotes number of substitutions occurring on average per site. Each clade has a corresponding pie chart detailing the proportion of elements that fall in each accessibility cluster, compared to all accessible ORR1Es (top right) (**E**) Plot of all motif and proto-motif matches for PU.1 on positive (top) or negative (bottom) strand among ERVL-MaLR subfamily consensus sequences present in Fig. 1D, which are significantly enriched in at least 4 cell types. (**F**) Percentage of ORR1Es among T, ILC3 and MF clusters with PU.1 motif minus percentage of background ORR1Es with PU.1 motif, relative to the ORR1E consensus sequence. Insert indicates percentage of ORR1Es in each cluster with a ROR TF motif at the dotted position in the ORR1E consensus.

### Origin of TF motifs within ODEs

The divergent accessibility patterns, TF motif enrichment and preferential TF binding across ODE loci suggest that sequence changes have shaped these elements as CREs to function across immune cells. Such mutations could have occurred in ancestral copies that then spread by transposition, or by point mutation in individual copies after their insertion, or a combination of both. The first model predicts that ODEs from each cluster would form distinct phylogenetic clades. To test this, we performed a phylogenetic analysis on accessible ORR1E (Fig. 3D) and ORR1D2 elements (Fig. S3D). Using a threshold of >90% bootstrap support [75] and at least 100 elements to define clades, we identified 5 clades among ORR1E and 2 among ORR1D2 elements. For each clade, we tallied the distribution of ODE loci that fell into each of the accessibility-derived clusters, revealing that each clade included an even distribution of loci from each cluster (Fig. 3D & S3D). These results do not support a model whereby each ODE accessibility cluster corresponds to the spread of ancestral copies with distinct combination of TF motifs. Rather, these results suggest that ODEs acquired different motif combinations via mutation after their insertion in the genome.

We next sought to understand why ODEs are enriched in accessibility across immune cell development while related LTR subfamilies that also expanded in rodents (ORR1B1, ORR1B2, ORR1F and MTD) are not. We hypothesized that ODEs had a greater propensity to evolve into CREs owing to a wider “starting set” of lineage-specifying TF motifs within their ancestral sequence. Thus, we searched for TF motifs among ODE and related TE LTR subfamily consensus sequences, which are a proxy for their ancestral sequence [76,77]. We categorized sequences as either perfect motif matches, or as proto-motifs when they were 1-2 nucleotides away from a perfect match. ORR1E was the only subfamily that harbored two PU.1 motifs and one PU.1 proto-motif, while other subfamilies contained at most a single PU.1 motif (Fig. 3E). The ORR1E consensus also contains a ROR-family motif (Fig. S3E), two E2A motifs (Fig. S3F) and two combined ETS-RUNX motifs (Fig. S3G), motifs largely absent from other ORR consensus sequences. Both ORR1E and ORR1D2 consensus sequences possess a unique RUNX-family motif (Fig. S3H) and proto-motifs for CTCF (Fig. S3I). These observations suggest that ODE ancestral sequences contained a diverse set of preexisting motifs for lineage-specifying TFs, imbuing an elevated potential to evolve *cis*-regulatory activity in a wider range of immune cell types than other related LTR subfamilies.

The presence of lineage-specific TF motifs in the ODE ancestral sequence does not fully explain the diversity of ODE accessibility and TF binding patterns across immune cells. We hypothesized that, post insertion, loci acquired diverse *cis*-regulatory activity across the immune system via 1) differential maintenance of ancestral TF motifs, and/or 2) acquisition of new motifs by mutations at proto-motif or *de novo* sites. To trace the origins of lineage-specifying TF motifs enriched across ODE clusters, we quantified the maintenance of ancestral motifs and their acquisition by mutation for ORR1E (Fig. S4A) and ORR1D2 (Fig. S4B). While these ancestral motifs are preferentially maintained across ODEs, the site(s) maintained differ across ODE clusters. For example, compared to background ORR1Es, “MF” ORR1Es preferentially maintain consensus PU.1 motifs at positions ∼120 and ∼170, whereas “T cell” ORR1Es tend to maintain the consensus PU.1 motif at ∼170 only. “ILC3” ORR1Es preferentially lose the consensus PU.1 motif at position ∼120 through mutations that strengthen the overlapping ROR-family motif (Fig. 3F). Thus, differential preservation of ancestral motifs contributed to diversification of ODE loci as CREs across cell types.

TFs that only had proto-motif(s) in the ODE consensus sequences can be split between two groups: 1) the majority preferentially acquired motifs at proto sites – CTCF and GATA, or 2) the motifs were acquired both at proto sites and *de novo* – EBF1, Tbet and Maz (Fig. S4A & S4B). In sum, subsets of ODEs acquired distinct TF motif repertoires both by differential preservation of ancestral motifs, and by the gain of new motifs either from proto-motifs or *de novo* through more extensive mutations (requiring >2 substitutions).

### ODEs likely regulate hundreds of cell type-specific genes across immune cell development

To explore the role of ODEs in regulating adjacent genes, we predicted CRE-gene pairs across immune cell development. We searched for a strong correlation in accessibility of distal ATAC-seq peaks (∼300,000) with gene expression across 85 cell types, using matched RNA-seq data [36]. To identify the strongest CRE-gene pairs, we calculated all possible Spearman correlation coefficients for each putative CRE-gene pair on the same chromosome. Out of ∼8.7 million pairs within 1Mb, we filtered for the strongest correlations for a given CRE to all genes on the same chromosome (Fig. 4A). This approach yielded 198,729 strongly correlated CRE-gene pair linkages (∼2.3%; Fig. S5A), of which 65,827 (∼33.1%) were TE-derived, including 2,226 ODE-derived pairs. ORR1E had 1,441 (∼2.6%; Fig. 4B) filtered pairs, while ORR1D2 had 785 (∼2.7%; Fig. S5B). CRE-gene correlations were either positive, indicative of a CRE activating gene expression, or negative, indicative of CRE-mediated repression.

**Fig. 4:**
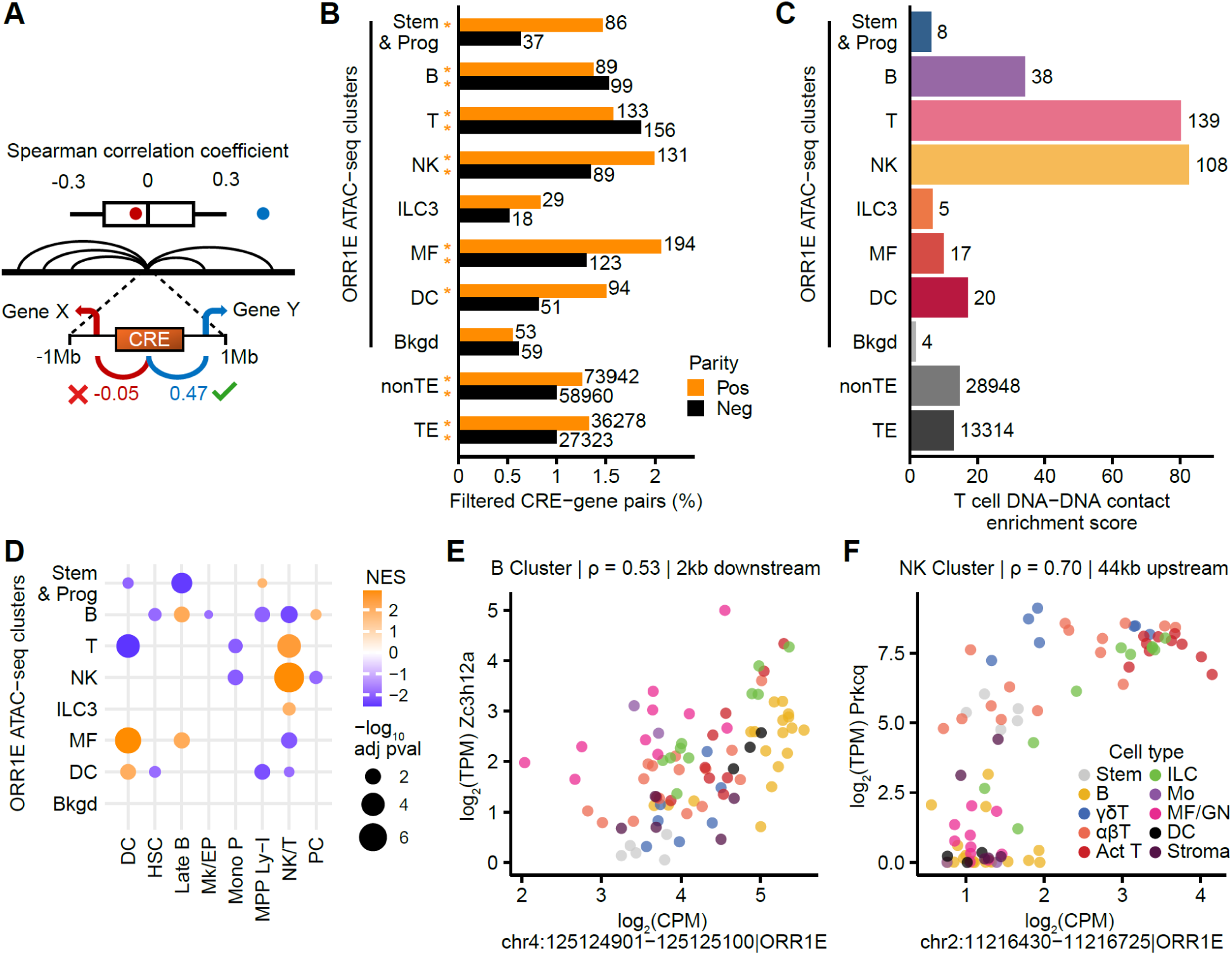
ODEs regulate hundreds of genes across immune cell development. (**A**) Cartoon representing Spearman correlation filtering process. After calculating all possible Spearman correlation coefficients between each individual peak’s (putative CRE) accessibility and every gene’s expression on the same chromosome, we retained CRE-gene linkages if their linear distance was <1Mb and their correlation coefficient was an outlier: either lower than the 25^th^ quartile - [1.5 * IQR] or higher than the 75^th^ quartile + [1.5 * IQR] for each CRE-specific distribution of correlation coefficients. (**B**) Barplot showing the proportion of filtered CRE-gene linkages for each ORR1E cluster and all other TE-derived or non-TE-derived peaks, split by positive (orange) and negative correlation coefficient (black) with orange asterisk indicating significantly (Fisher’s Exact Test adjusted p-value<0.05) higher proportion of filtered interactions than the respective ORR1E background cluster. (**C**) Barplot showing enrichment of DNA-DNA contacts in CD8+ T cells per ORR1E cluster, compared to all non-TE and TE peaks. (**D**) Dotplot showing significant (adjusted p-value<0.05) GSEA results of ORR1E gene targets for cell type marker gene sets [82]. (**E – F**) Scatterplots showing ATAC-seq and RNA-seq values for a single ORR1E-gene pair with supporting DNA-DNA contacts. Highlighted genes are leading edges from cluster GSEA enrichments in Fig. 4D, with Zc3h12a from the Late B gene set (E) and Prkcq from the NK/T gene set (F).

To determine whether ODEs with cell type-specific accessibility served as likely regulators of nearby genes, we compared the fraction of filtered CRE-pairs across ODE clusters to “background” (Fig. 4B & S5B) and found that all ODE clusters except “ILC3” had significantly more positive pairs compared to “background” ODEs. Thus, cell type-specific ODEs are more likely to regulate nearby genes compared to “background” ODEs. To investigate whether ODEs were more likely to serve as CREs relative to accessible sequences genome-wide, we compared the proportions of filtered pairs (both positive and negative) in ODE clusters against all other TE-derived and non-TE-derived peaks. We found that “B”, “T”, “NK”, and “MF” ODE clusters all had significantly higher proportions of filtered pairs (Fig. S4C & S4D), implicating that a subset of ODEs have been co-opted as CREs across immune cell development.

To validate predicted ODE-gene regulatory interactions, we performed Micro-C [78] in mouse naïve CD8+ T cells to capture CRE-promoter contacts. To identify DNA-DNA contacts associated with CREs, we used the Activity by Contact (ABC) model [79], which uses contact information, chromatin accessibility and H3K27ac ChIP-seq signal to infer contacts between promoters and candidate CREs. In addition to our Micro-C data [80], we used H3K27ac ChIP-seq [81] and ATAC–seq data [36] to predict 42,782 CRE-gene interactions in CD8+ T cells. In total, 4,137 (∼9.7%) interactions predicted by the ABC model were also identified as CRE-gene pairs in our ATAC-RNA correlation analysis, a significant overlap (Fisher’s Exact Test, p-adjust<0.05). Overall, the ABC model predicted 339 ORR1E-gene interactions and 181 ORR1D2-gene interactions in CD8+ T cells.

To assess whether ODEs were overrepresented in CRE-gene interactions predicted by the ABC model, we calculated an enrichment score for each ATAC-seq cluster for ORR1E (Fig. 4C) and ORR1D2 (Fig. S5E), and all other non-TE or TE-derived peaks. This enrichment score measures overrepresentation of peak-gene contacts for each set relative to their genomic proportion. ODE-gene interactions were overrepresented amongst predicted CRE-gene pairs, with 6-83 fold enrichment for ORR1E (Fig. 4C) and 10-74 fold for ORR1D2 (Fig. S5E), depending on the ODE cluster identity. “T” and “NK” clusters had the strongest enrichment scores for both ORR1E and ORR1D2, concordant with their similar developmental trajectory and function. In contrast, background clusters had the smallest enrichment scores (∼2 fold), consistent with their non-specific accessibility patterns, decreased propensity for TF binding and decreased number of filtered gene pairs in the RNA-ATAC correlation analysis. Together, these results implicate ∼2,200 ODE loci as CREs throughout immune cell development with 272 ODE loci contacting 506 gene promoters in T cells.

### Predicted ODE-regulated genes are enriched for those that define cell type identity

We sought to determine whether genes predicted to be regulated by ODEs were relevant to the biological identity of each cell type. We considered ODE-gene interactions identified by either the correlation analysis or ABC model (n=2,698) and performed gene set enrichment analyses (GSEA) across each ORR1E (Fig. 4D) and ORR1D2 cluster (Fig. S5F) for genes that mark cell types throughout early immune cell development [82]. All ORR1E clusters, except background, had at least one significant enrichment; and strikingly, these enrichments were consistent with the identity of the respective cell type(s). For example, ORR1E “T”, “NK” and “ILC3” clusters had strong enrichments for NK/T associated gene sets, while “MF” and “DC” clusters had strong enrichments for DC gene sets. Furthermore, the “B”, “MF” and “DC” clusters were depleted for gene sets associated with NK/T cells, concordant with immune cell development requiring repression of genes specifying other lineages [83,84]. In particular, key lineage-specifying TFs can repress differentiation towards a particular lineage, such as Pax5 in B cells inhibiting T cell development and GATA-3 in T cells repressing B cell development [85]. We observed similar trends for ORR1D2 (Fig. S5F). These results suggest that predicted ODE-gene linkages are associated with gene sets contributing to cell type identity.

To further understand the biological relevance of ODE regulatory interactions, we investigated the function of ODE-gene pairs that contributed most to enrichment of cell type identity and had ODE-promoter contacts in naïve CD8+ T cells. Out of 27 genes (22 for ORR1E, 4 for ORR1D2, 1 linked to both), nearly all (20) had established roles in their respective cell types (Table S4). For example, *Regnase-1* (Zc3h12a) is necessary for B cell response [86] and activation of T cells and macrophages [87], and its expression is correlated (ρ = 0.53) with the accessibility of its ODE pair across immune cell development (Fig. 4E). This ORR1E locus is located ∼2 kb downstream of the *Regnase-1* promoter, physically contacts the promoter, and is bound by Pax5 in B cells, Runx1 in T cells and PU.1 in macrophages, concordant with *Regnase-1*’s functions in these cells (Fig. S5G). We also identified a notable correlation (ρ = 0.70) between an ODE and PKCθ (Fig. 4F), a kinase preferentially expressed in T cells and necessary for T cell activation [88]. These data suggest that many ODEs have been co-opted as CREs acting on genes defining specific immune cell identities.

### ODEs drive rodent-specific changes in gene expression

Given that ODEs are rodent-specific, we investigated whether ODEs contribute to changes in gene expression in mouse relative to human. We considered two models: 1) the “booster” model whereby ODE-derived enhancers drove rodent-specific increases in expression relative to species lacking these elements, such as humans, and/or 2) the “recruitment” model whereby ODEs introduced enhancers that recruited genes into a cell type where they were not previously expressed. To investigate the “booster” model, we compared expression of ODE-regulated genes in mouse with their ODE-lacking human orthologs across homologous cell types. We utilized expression data for 12 different human immune cells [89] and calculated a scaled expression score for orthologs. Genes predicted to be under control of an ODE enhancer were significantly more highly expressed in mouse than in human. This trend extends to nearly all cell types investigated across ODEs, with 10/12 cell types for ORR1E (Fig. 5A-B & S6A-J) and 10/12 for ORR1D2 (Fig. S6K-V). These results suggest that ODE-derived enhancers frequently act as boosters of cell identity.

**Fig. 5:**
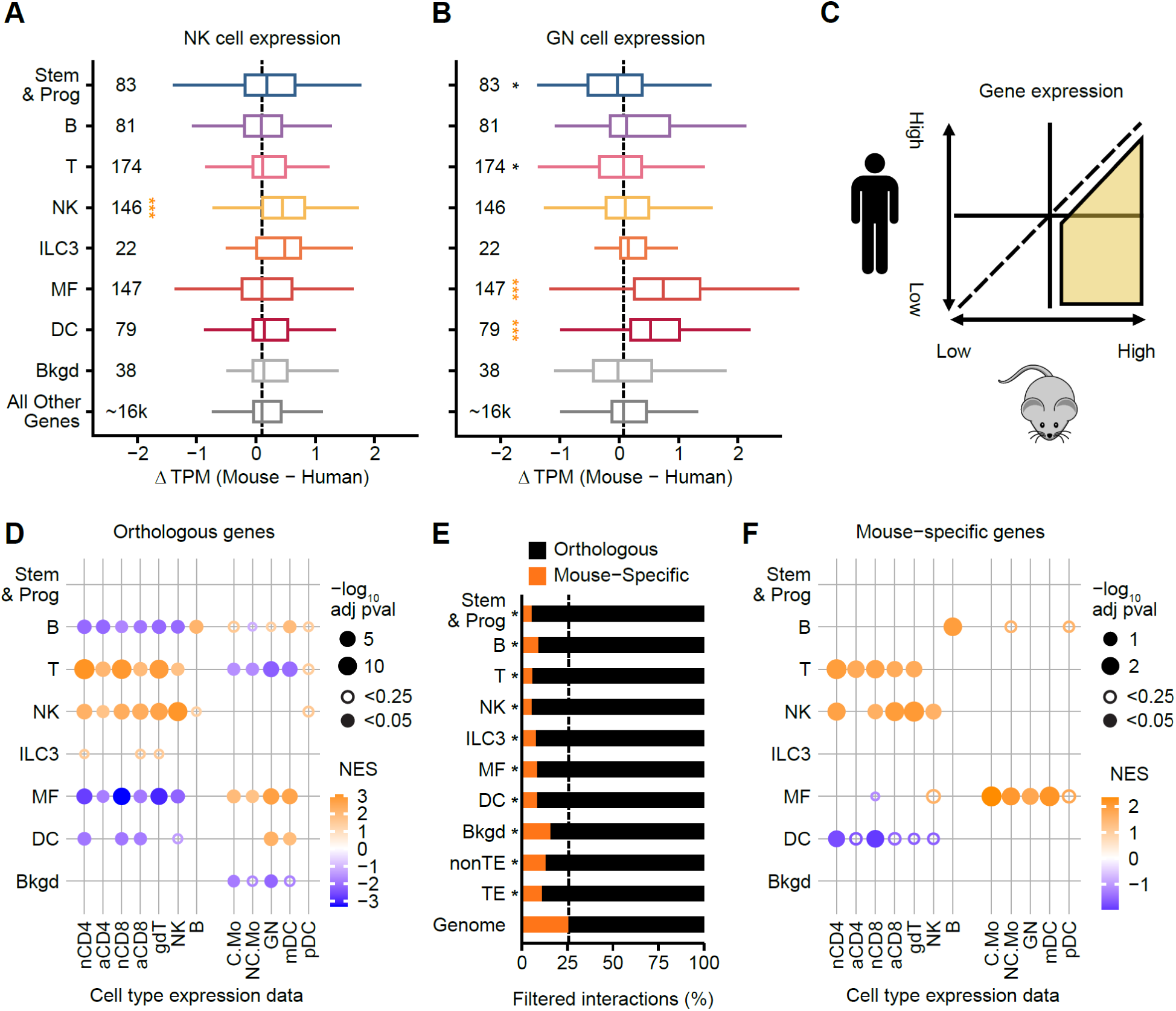
ODE-linked genes have mouse cell type-specific increases in expression compared to human. (**A – B**) Boxplots of positively correlated gene targets across each ORR1E cluster and all other genes measuring the significant (Wilcoxon Rank Sum Test compared to “All Other Genes”, adjusted p-value<0.05; black or orange asterisks indicate decreased or increased expression, respectively) distribution of scaled difference in gene expression between mouse and human in Natural Killer cells (NK, panel A) or Granulocytes (GN, panel B), indicating the numbers of genes per group. *p<0.05, ***p<0.0005. (**C**) Cartoon representing process for selecting orthologous genes between mouse and human to use in gene set enrichment for D. (**D**) Dotplot showing significant (adjusted p-value<0.25 is open circle, adjusted p-value<0.05 is filled circle) GSEA results of orthologous ORR1E gene targets for gene sets that are more expressed in mouse compared to human. (**E**) Barplot of proportions of CRE-gene linkages to genes orthologous between mouse and human (black) or mouse-specific (orange). Black asterisks indicate significantly (Fisher’s Exact Test adjusted p-value<0.05) lower proportions of mouse-specific target genes compared to genomic proportion. (**F**) As per panel (D) but using mouse-specific ORR1E gene targets.

Taking NK cells as an example, we found that genes regulated by “NK” ODEs were the only set of genes that had significantly increased expression in mouse compared to human (Fig. 5A). Notably, cell types such as granulocytes had “Stem & Prog” and “T” ORR1E-regulated genes significantly decreasing in expression compared to human alongside significant increases in expression of “MF” and “DC” ORR1E-regulated genes (Fig. 5B). Conversely, genes predicted to be negatively regulated by ODEs were significantly less expressed in mouse compared to human in 4/12 cell types for ORR1E and 2/12 cell types for ORR1D2 (Fig. S7). We next determined whether ODE-target genes were enriched for orthologous genes more expressed in mouse compared to human (Fig. 5C). Across ORR1E (Fig. 5D) and ORR1D2 (Fig. S8A) clusters, target genes were significantly enriched in gene sets concordant with accessibility patterns, such as “T” and “NK” ODE cluster target genes being positively enriched in genes highly expressed in T and NK cells. These “T” and “NK” ODE cluster target genes were negatively enriched in B, MF and DC cells, suggesting ODEs broadly repress T and NK relevant genes in non-cognate cells to reinforce cell type identity. Together, these results imply that introduction of ODEs in the rodent lineage altered gene expression levels in ways that reinforced cell type identities across the immune system.

To examine whether ODEs recruited new genes into specific regulatory networks, we assessed whether ODEs preferentially regulate mouse-specific genes. We defined 6,890 mouse-specific genes expressed across immune cells using orthology databases [90] and found that the majority (n=4,694; ∼68%) were not annotated as protein-coding. The proportion of mouse-specific genes per ODE cluster ranged from ∼5-16% for ORR1E (Fig. 5E) and ∼4-15% for ORR1D2 (Fig. S8B) compared to ∼24% in the genome; all significant depletions. Nevertheless, ODEs are predicted to regulate 169 mouse-specific genes (115 for ORR1E; 61 for ORR1D2). Notably, mouse-specific ORR1E-target genes are significantly enriched in highly expressed cell type-specific genes (Fig. 5F), with mouse-specific “MF & DC” ORR1D2-target genes significantly enriched for highly expressed monocyte genes (Fig. S8C). These results suggest that ODEs reinforce immune cell type-identity, both by boosting immune cell type-specific gene expression in mouse and by incorporating mouse-specific genes into cell type-specific regulatory networks.

### ODE-derived CREs are under functional constraint

To further validate that ODEs have been co-opted for regulatory functions, we examined their sequence constraints across rodents. We predicted that if ODEs with cell type-specific accessibility were co-opted as CREs, they would show greater sequence conservation than “background” ODEs, indicative of purifying selection. We used phastCons [91] to measure evolutionary constraint across rodent species containing ODEs [92]. We used full length non-accessible ODEs (“full-length background” ODEs) as a set of elements most likely to evolve neutrally [37,93]. We found that ORR1E (Fig. 6A) and ORR1D2 (Fig. S8D) loci from the “B”, “T” and “NK” clusters had significantly higher phastCons scores than “full-length background”; ORR1E from the “MF” cluster were also significantly more conserved. All other cell type-specific accessible ODEs manifested higher average phastCons scores, although these individual trends were not statistically significant. These results suggest that a subset of ODEs with cell type-specific accessibility harbor sequences that are more evolutionary constrained than other ODEs.

**Fig. 6:**
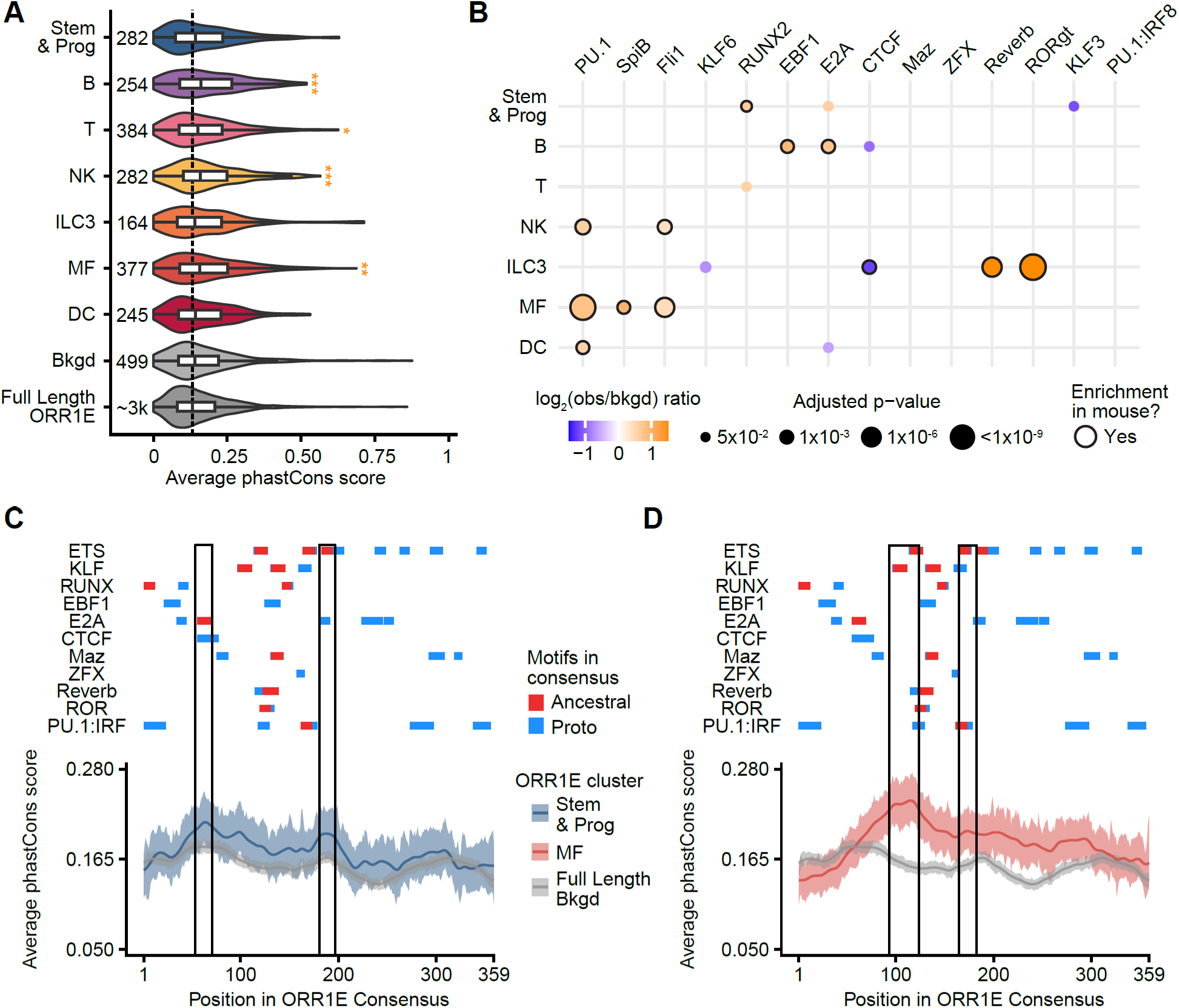
Accessible ODEs have evidence of selection across rodents. (**A**) Violin plot measuring significantly (Wilcoxon Rank Sum Test, adjusted p-value<0.05) higher average phastCons scores for ORR1E clusters compared to full-length non-accessible ORR1Es. Numbers indicate loci included per cluster. *p<0.05, **p<0.005, ***p<0.0005. (**B**) Dotplot of significant (binomial test adjusted p-value<0.05) motif enrichment results from HOMER on orthologous ORR1E loci in rat for each cluster compared to the “background” cluster. Black outline signifies if that cluster’s motif enrichment is present in the orthologous mouse cluster. (**C – D**) Visualization of matched motifs (Ancestral, red) or proto-motifs (Proto, blue) in ORR1E consensus for enriched motif clusters from Fig. 2B. Density plots with 95% confidence interval of average phastCons scores for ORR1E “Stem & Prog” cluster (C) or ORR1E “MF” cluster (D) and full-length background ORR1Es aligned to the consensus sequence. Boxed regions are representative local maxima of average phastCons scores.

The signal of purifying selection detected through phastCons analysis suggests that ODE activity as CREs preceded the divergence of mouse from other rodent species, such as rat, which diverged ∼12-24 million years ago from mouse [94]. This model predicts that ODEs with conserved *cis*-regulatory activity in mouse and rat share ancestral TF binding sites. Alternatively, they could have acquired TF binding sites independently in each lineage. To investigate these possibilities, we identified mouse ODEs with a rat ortholog and performed TF motif enrichment analysis for each rat ODE cluster against orthologous “background” ODEs. We found that rat ORR1E (Fig. 6B) and ORR1D2 (Fig. S8E) clusters are enriched for motifs shared with those in mouse (Fig. 2B & S2D). Shared enrichments include both ancestral motifs such as PU.1, and RORgt in the orthologous ORR1E ILC3 cluster. Depletion (indicative of loss) of the CTCF motif in ORR1E ILC3 clusters was also found in rat ODEs. Interestingly, orthologous ODEs contained rat-specific motif enrichments such as KLF6 in ORR1D2 “B” and “NK” clusters (Fig. S8E) as well as depletions such as KLF6 in ORR1E “ILC3” (Fig. 6B) and EBF1 in ORR1D2 “Stem & Prog” clusters (Fig. S8E). These observations suggest that ODEs in mouse and rat share similar cell type-specific TF motifs, including those present ancestrally upon ODE insertion and those acquired by mutation post genomic insertion. Thus, ODEs display a mixture of conserved and more recently evolved motifs.

We next sought to understand which TF motif(s) had the highest signature of purifying selection per ODE cluster. We visualized the average phastCons score across each cluster, overlaying motif matches and proto-motifs for enriched motifs for each ORR1E (Fig. S9A-F) and ORR1D2 (Fig. S10A-F) cluster. Each cluster had distinct profiles of high-scoring motifs, such as an E2A motif for the ORR1E “Stem & Prog” cluster (Fig. 6C). E2A promotes developmental progression of early immune cell progenitors [95] and is essential for commitment to B [96] and T cell [97] lineages. By contrast, ETS and KLF family motifs are the most conserved sequences for the ORR1E “MF” cluster (Fig. 6D), consistent with roles of ETS and KLF TFs in macrophages [98,99]. Together, these results suggest that each ODE cluster has a distinct set of TF motifs under purifying selection, enabling their cell type-specific regulatory activity.

## Discussion

The immune system consists of numerous cell types with distinct regulatory and functional niches, each specified by distinct regulatory networks governing their developmental trajectory and unique identity. Immune cell regulatory networks vary across vertebrates, leading to different proportions and phenotypes of immune cells and species-specific gene expression programs [100]. How these cell type-specific regulatory networks arose and how networks are differentially modulated across species remain poorly understood. Here, we show that two closely related TE subfamilies – ORR1D2 and ORR1E (ODE) – have been a pervasive source of CREs fueling the evolution of the rodent immune system. Through their expansion early in rodent evolution, ODEs have dispersed a vast reservoir of latent CREs, a subset of which has been co-opted to wire lineage-specific gene regulatory networks dictating immune cell identities. Our study indicates that thousands of ODEs display cell type-specific signatures of *cis*-regulatory activity throughout immune cell development. In fact, ODEs with *cis*-regulatory activity comprise only ∼0.03% of the genome but contribute ∼1.2% of all predicted CREs across mouse immune cell types. We show that a subset of ODEs with CRE activity have evolved under functional constraint since their insertion in the genome of a rodent ancestor, and that they contribute to divergent expression programs between mouse and human. This study adds to a growing body of evidence pinpointing TEs as a major source of regulatory innovation in mammalian genomes [23–26,28–35,101].

Prior studies have shown that TEs are an abundant source of species-specific regulatory elements. For example, TEs have been implicated in lineage-specific rewiring of the regulatory network of embryonic stem cells [102] and as the source for the majority of primate-specific TFBS and *cis*-regulatory elements [13,103]. However, it has not been shown previously that a single TE family serves as cell type-specific CREs across an entire developmental cascade, such as ODEs across the mouse immune system. Our study deepens understanding of TE co-option as CREs, through both the preservation and gain of TF binding sites, highlighting TEs as sources of species-specific regulatory novelty.

One of the most unexpected findings of this study is the partitioning of CRE activity of ODEs across diverse immune cell types. Previous studies have found that such CRE diversification within the same TE family typically results from the split and spread of different subfamilies, each with a unique combination of TF binding sites and expression patterns. For example, retroviral LTRs frequently undergo regulatory diversification to form new subfamilies with different expression niches [46,47,104]. This diversification process has been interpreted as a form of partitioning potentially driven by competition between subfamilies and/or escape from genome defense systems [46,105,106]. However, we found that diversification of ODEs as CREs largely occurred after their insertion in the genome as cell type-specific accessibility patterns do not stratify across phylogenetic clades. Indeed, we show that ODEs likely functionalized as cell type-specific CREs through selective maintenance of ancestral TF motifs and acquisition of novel TF motifs through mutation post insertion. This interpretation is akin to both subfunctionalization (splitting of functions between copies) and neofunctionalization (acquisition of new functions) models of gene duplication evolution [107]. Previous studies have shown that

TEs can repeatedly acquire and/or remodel TF motifs through mutation post-insertion, such as Alu elements via CpG deamination [108,109], RSINE1 elements morphing into circadian enhancers [110], or a MER20/MER39 composite element that acquired *cis*-regulatory activity to regulate decidual prolactin in human pregnancy [111]. Our findings expand on these studies by describing ODEs as cell type-specific CREs across mouse immune cell development, with diverse regulatory activities acquired post-insertion through functionalization.

Our cross-species analyses implicate ODEs as rodent-specific “boosters” of cell type-specific gene expression across immune cell development. We correlated ODE infiltration of the rodent lineage to the upregulation of hundreds of relevant genes and to the recruitment of dozens of genes into established cell regulatory networks, including mouse-specific genes. Such lineage-specific “boosting” by TE-derived enhancers is consistent with studies identifying TEs as key sources of species-specific and context-specific regulatory novelty [3,26,35,112]. One example is LTR5HS, the LTR of ape-specific HERVK insertions, which serve as enhancers in human preimplantation embryos but not in rhesus macaque, which lacks LTR5HS [113,114]. These observations beg the question: what makes ODEs special compared to other TEs?

We propose that ODE’s capacity for broad *cis*-regulatory activity in immune cell development has been driven by 1) the presence of ancestral TF binding sites for key immune cell development regulators, such as PU.1, 2) the abundance of these elements in the genome (∼43k loci) and 3) their relatively old age, entailing ∼70 million years of evolution [92], providing opportunity to diversify their cis-regulatory activity by mutation. As ODEs are distinct for the presence of TF motifs relevant to immune cells, it is intriguing to consider that their expansion may have been tolerated due to their intrinsic ability to serve as batteries driving rodent immune evolution.

ODE-target genes include necessary components of T cell activation and immune response pathways, suggesting that ODEs are serving a critical role in shaping rodent immune cell evolution. While our results show that ODE-target genes have cell type-specific changes in gene expression compared to cognate human cells, it is unclear whether ODEs regulate the same or even different genes in other rodents. The ODE-driven modulation of genes necessary for immune function and development across rodents may have facilitated species-specific protection against rapidly evolving pathogens. Further investigations into the accessibility patterns, TF binding profiles and phenotypic impact of ODEs in other rodents across immune cell development are necessary to understand when ODE-derived co-option occurred, either rapidly upon insertion in the rodent ancestor, and/or over evolutionary time functionalizing as cell type- and species-specific CREs. Finally, investigating the propensity of other TEs to functionalize as CREs across other developmental cascades or rapidly evolving systems will help elucidate the principles underlying the creation of cell type-specific CREs and the incorporation of new genes into regulatory networks.

## Methods

### Annotation of TEs

TEs were annotated using the Dfam release 3.6 [76] as input to RepeatMasker version 4.1.2-p1 (http://www.repeatmasker.org)

### Annotation of TE-derived ATAC-seq peaks

Peaks within each sample were annotated as being TE-derived if the summit overlapped an annotated TE. For unified peaks, peak summits across all samples were intersected against TE annotations and then against unified peaks. If a unified peak contained at least one summit that intersected a TE, it was annotated as TE-derived, and non-TE-derived otherwise.

### ATAC-seq peak calling

Raw fastq files were downloaded from GEO using fasterq-dump version 3.1.1 (https://github.com/ncbi/sra-tools). Paired-end reads were trimmed using Trim Galore version 0.6.5 (https://github.com/FelixKrueger/TrimGalore), aligned to the mm10 genome assembly using STAR version 2.7.5a [115], deduplicated with Picard MarkDuplicates (https://github.com/broadinstitute/picard), and filtered against the ENCODE blacklist for mm10 [116] and regions with high sequence similarity to the mitochondrial genome. Peaks were called across each sample by combining replicates and using MACS3 [117] with parameters “-f BED -g mm -q 0.001 --nomodel --shift -75 --extsize 150 -B --keep-dup all --SPMR”. A unified peak set was generated by first expanding the summits of each sample by the median peak size in that sample and then merging using bedtools [118]. Counts for the unified peak set were calculated by intersecting the paired fragments from each sample and then normalized via quantile normalization using qnorm (https://pypi.org/project/qnorm/).

### Calculation of expression difference between mouse and human genes

RNA-seq data from ImmGen [36] and the Human Protein Atlas [89] were separately normalized via quantile normalization using qnorm (https://pypi.org/project/qnorm/) and then scaled and centered in R to generate a relative measure of expression for each gene within mouse and human. The relative expression values of orthologous human genes were then subtracted from the expression of their orthologous mouse gene(s) for each matched cell type to calculate relative expression difference between mouse and human.

### Correlation and filtering of peak-gene linkages

Using the Refseq Curated [119] gene set, each of the unified ATAC peaks (putative CREs) were paired with all genes on the same chromosome. We calculated the Spearman correlation between the accessibility and expression of each CRE-gene pair. For each CRE, we utilized the total distribution of Spearman correlation scores to identify outliers, which had scores that exceeded (third quartile + (IQR * 1.5)) or fell below (first quartile – (IQR * 1.5)) the overall distribution. All outlier CRE-gene pairs were annotated as filtered and non-filtered otherwise.

### DNA-DNA contact enrichment analysis

We calculated the enrichment of DNA-DNA contacts called by the ABC model [79] in T cells as follows:

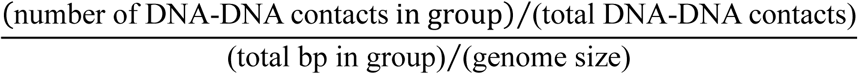

### Identification of orthologous and species-specific genes between mouse and human

We used the R package orthogene (https://doi.org/doi:10.18129/B9.bioc.orthogene) to access the babelgene (https://igordot.github.io/babelgene/), g:profiler [90], and homologene [120] orthology databases. We combined the hits from all databases for orthologous genes between mouse and human and filtered for genes with expression data in both species. Genes with no hits from one species to the other were annotated as species-specific.

### Permutation shuffle test

For each accessibility dataset, we performed 1,000 iterations, by randomly shuffling the summit for each peak to a new position on the same chromosome, maintaining the distance to the nearest TSS. These shuffled positions generated an expectation of overlap with accessibility. For each iteration, all subfamilies were measured for enrichment, as determined by at least 2-fold more observed summit overlaps compared to shuffled summits, and depletion via at least 2-fold fewer observed summit overlaps compared to shuffled summits. The resulting permutation p-value was calculated as:

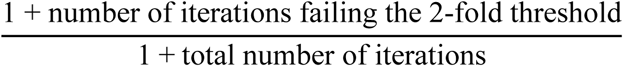

Thus, a p-value with a numerator of 1 had all iterations return at least 2-fold more/fewer observed summit overlaps than expected. Finally, the number of expected summit overlaps was averaged across all iterations and used as input to a Fisher’s Exact Test to measure significant enrichment and depletion, and the resulting p-value was adjusted for multiple tests across each dataset using the Benjamini & Hochberg method [121].

### PhastCons analyses

We intersected ODEs against the glire subset of the mouse 60-way phastCons dataset from the UCSC genome browser [91,122]. The output of phastCons is a base-wise value ranging from 0-1, where a base scoring 0 is likely to have evolved under neutrality while a base scoring 1 is likely to be constrained by purifying selection. Average phastCons scores were calculated for each ODE, whereas alignments between individual ODEs and their respective consensus were used to assign base-wise phastCons scores across ODE clusters.

### Phylogenetic analyses

We aligned all accessible ORR1Es or ORR1D2s using MAFFT [123], trimmed uninformative positions using trimAl [124], and then created an unrooted approximately-maximum-likelihood phylogenetic tree using FastTree [125,126]. Clades were identified using the following criteria: 1) does a node contain >= 0.90 local support? 2) does the node and corresponding child nodes contain at least 100 elements? and 3) are there no nodes that match criteria 1 & 2 within the child nodes?

### Principle component analysis

Using the DESeq2 R package [127,128], PCAs were generated by performing a variance stabilizing transformation (vst) on the unified peak set counts, followed by the function plotPCA with the top one thousand most variable TE- or non-TE-derived peaks.

### Statistics and Reproducibility

All data points were included in statistical analysis. Statical analyses and plots were generated using R version 4.1.3.

## Data availability

All accession codes or download links for publicly available data are listed in Supplementary Data 1.

## Code availability

All code for analysis is available at https://github.com/Grimson-Lab/ODE_manuscript.

## Acknowledgments

We thank past and current members of the Feschotte and Grimson labs for their continuous support and caring community during lab meetings and social events. Special thanks go to Connor Kean and Michelle Liu.

## Author Contributions

J.D.C. analyzed and interpreted data, generated the figures, and wrote the manuscript. C.F. and A.G conceptualized the study, analyzed and interpreted data, and wrote the manuscript.

## Corresponding authors

Direct correspondence to Cédric Feschotte and/or Andrew Grimson

---

**Fig. S1:**
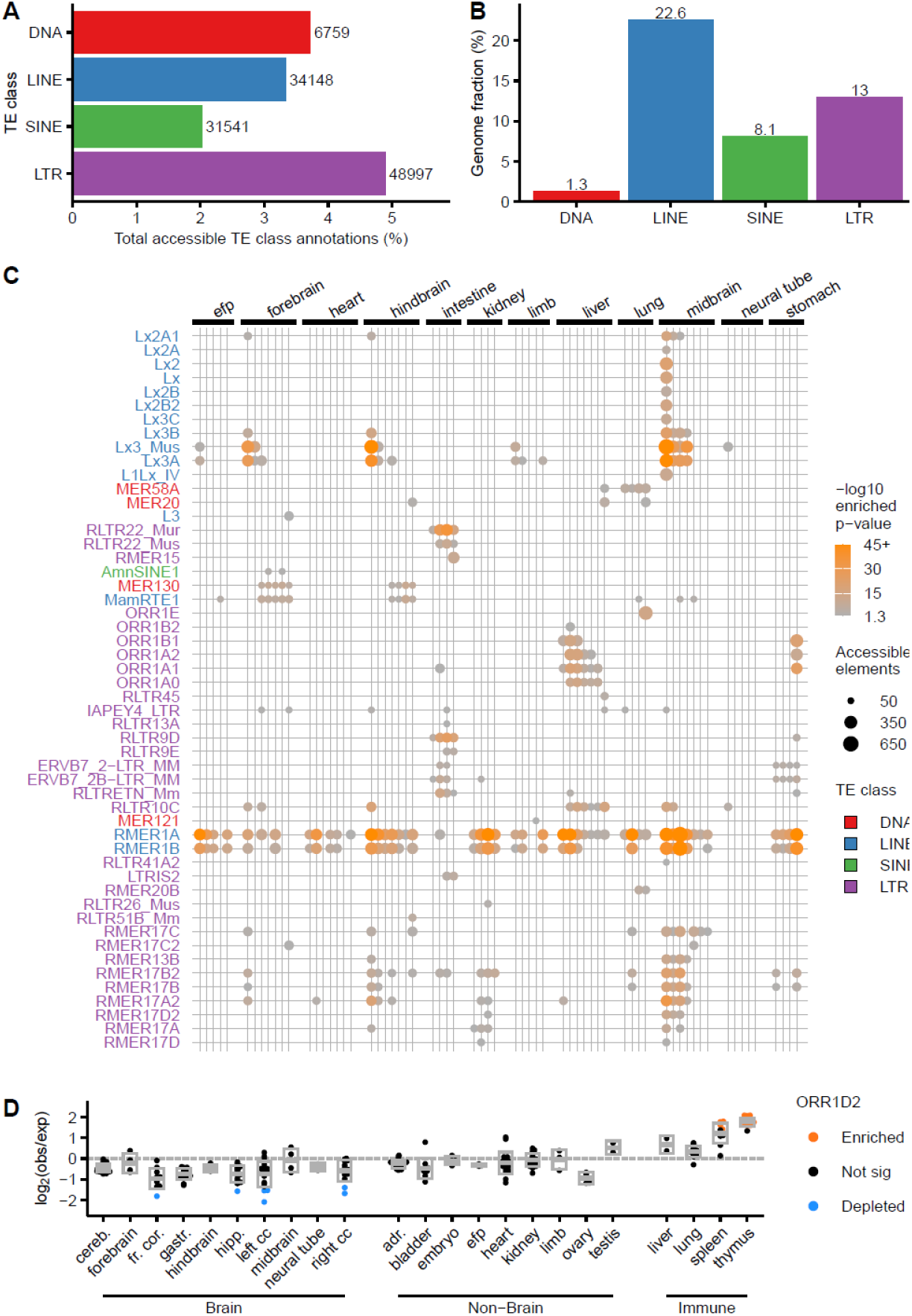
ODE accessibility is immune-cell specific. (**A**) Barplot showing percentage of TE classes that are accessible at least once across all Immgen [36] ATAC-seq datasets with numbers indicating number of accessible loci. (**B**) Barplot showing genomic fraction (in base pairs) of each TE class with numbers indicating rounded percentage. (**C**) Dotplot showing enrichment of TE subfamilies in accessibility, as in Fig. 1D, but using ATAC-seq data from ENCODE’s mouse developmental matrix. (**D**) Enrichment in accessibility of ORR1D2 across ENCODE’s mouse developmental matrix DNase-seq data as in Fig. 1E, where each dot represents a dataset taken at different developmental timepoints. Orange and blue dots indicate significant (permutation test<0.05 and Fisher’s Exact Test adjusted p-value<0.05) enrichment or depletion, respectively.

**Fig. S2:**
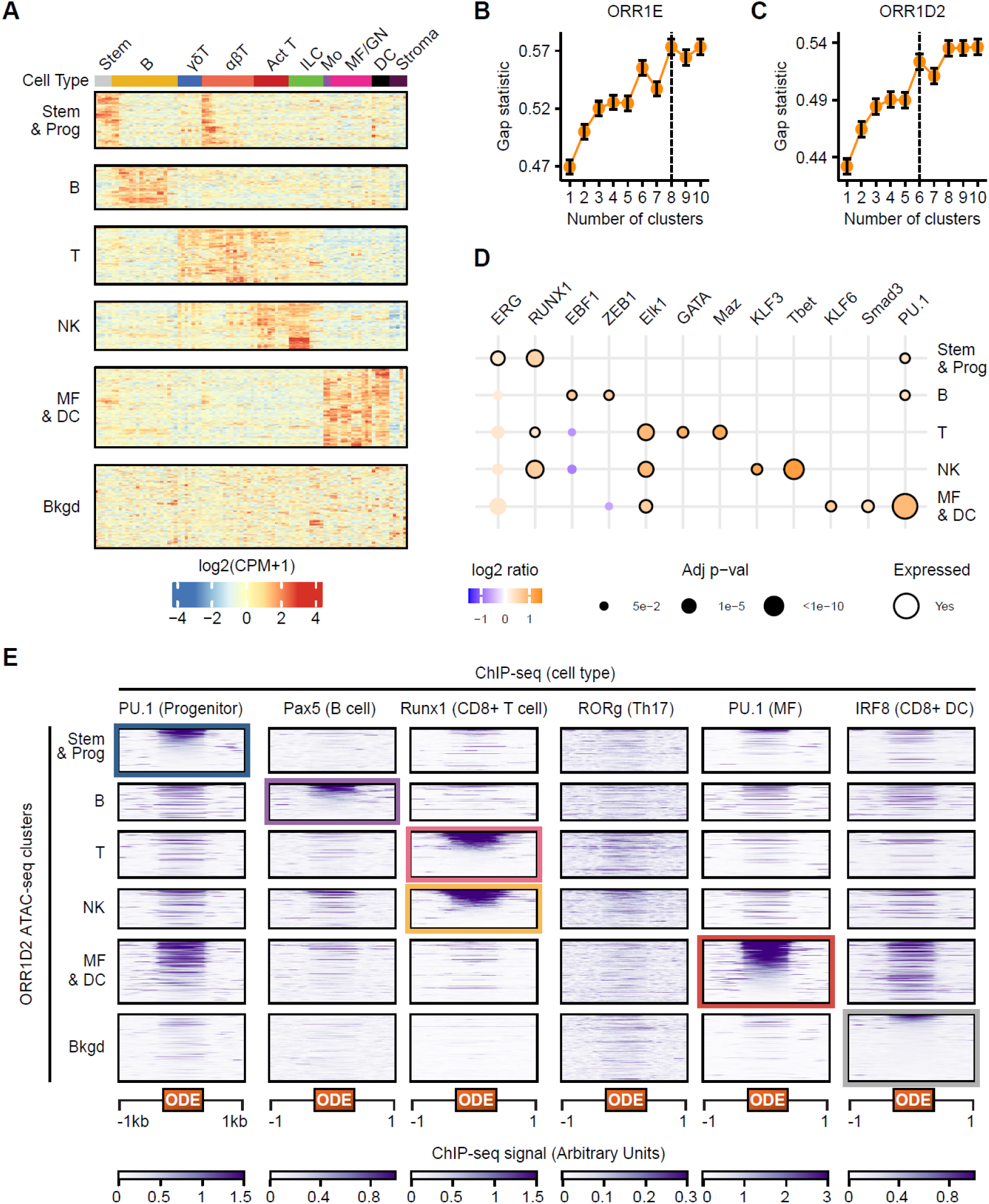
ORR1D2s stratify by accessibility patterns, TF motif enrichment and TF binding. (**A**) Heatmap of row-scaled quantile-normalized log_2_(CPM) ATAC-seq from Immgen across accessible ORR1D2 loci, plotted using unsupervised kmeans clustering. (**B – C**) Gap statistics for ORR1E (B) and ORR1D2 (C) to determine optimal number of clusters for kmeans clustering in Fig. 2A and Fig. S2A, respectively. (**D**) Significant (Binomial test adjusted p-value<0.05) motif enrichment results from HOMER on ORR1D2 clusters using cluster 6 (“Bkgd”) as background. Dots have black outline if they are expressed across majority of cell types driving accessibility clusters. (**E**) Heatmaps of TF ChIP-seq datasets across ORR1D2 clusters expanded by 1kb upstream and downstream of individual ORR1D2 loci for each accessibility cluster. Highlighted boxes indicate dataset used for descending row sorting by signal intensity for each cluster across all ChIP-seq heatmaps.

**Fig. S3:**
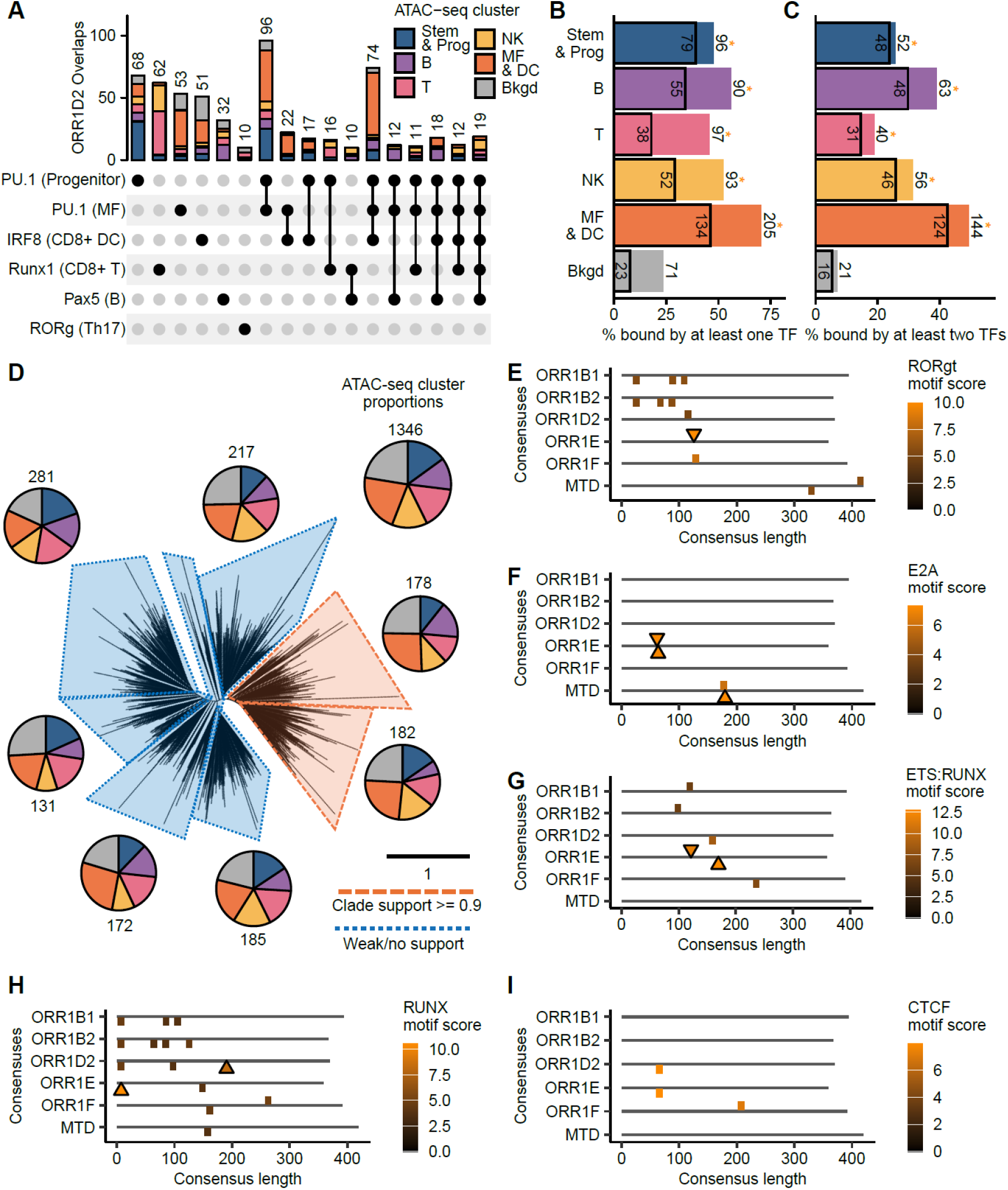
ODEs contain distinct TF repertoires compared to related subfamilies. (**A**) Upset plot, as in Fig. 3A, showing combinations of TF ChIP-seq binding patterns across ORR1D2 loci. (**B – C**) Barplots showing total proportion of loci in each ORR1D2 cluster that are bound by at least one (B) or two (C) TFs in distinct cell types. Box indicates proportion bound by PU.1 and * indicates significantly (Fisher’s Exact Test adjusted p-value<0.05) higher total % bound compared to “Bkgd” for non-boxed proportions. (**D**) Phylogenetic tree of accessible ORR1D2s highlighting presence of clade support (>= 0.90) in orange or weak/no robust clade support (< 0.90) and black scale bar denotes number of substitutions occurring on average per site. Each clade has a corresponding pie chart detailing the proportion of elements that fall in each accessibility cluster, compared to all accessible ORR1D2s (top right). (**E – I**) Plot of all motif and proto-motif matches for RORgt (E), E2A (F), ETS:RUNX (G), RUNX (H), and CTCF (I) on positive (top) or negative (bottom) strand among ERVL-MaLR subfamily consensus sequences present in Fig. 1D that are significantly enriched in at least 4 cell types.

**Fig. S4:**
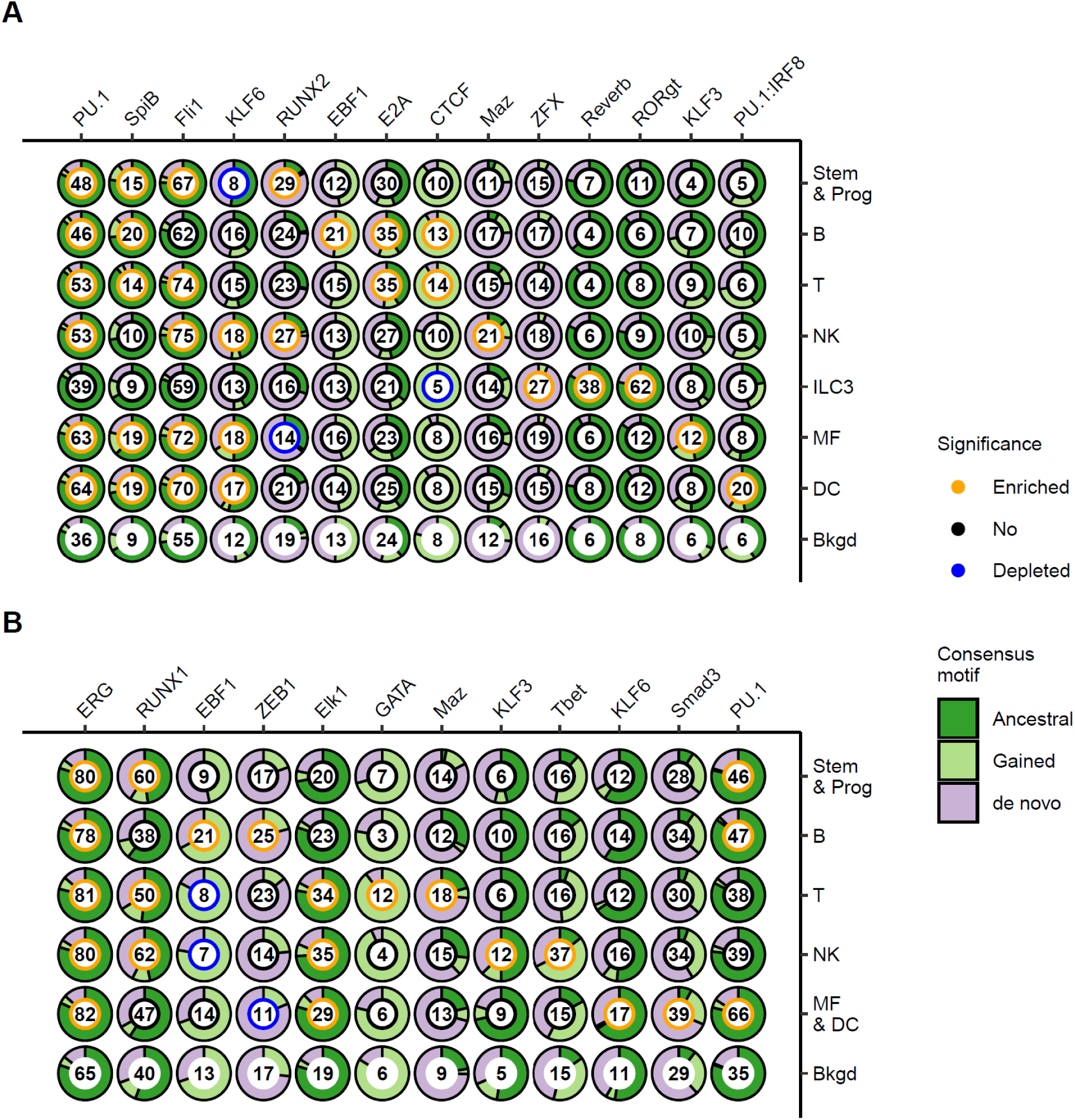
ODE loci variably maintained, gained or acquired *de novo* TF motifs. (**A – B**) Donut plots showing assigned motif origin for ORR1E (A) and ORR1D2 (B) loci across each cluster for all enriched motifs shown in Fig. 2B and Fig. S2D, respectively. Number within each donut is the rounded percentage of loci with at least one of the corresponding motifs, while the inner color indicates whether that motif is enriched (orange), depleted (blue), or not significantly different (black) compared to background, as in Fig. 2B and Fig. S2D. Dark green indicates the motif aligns to the ancestral motif in consensus (indicative of maintenance), light green indicates the motif aligns to an ancestral proto-motif (indicative of gain), while pink indicates the motif does not align to any prior ancestral motif (indicative of *de novo* acquisition).

**Fig. S5:**
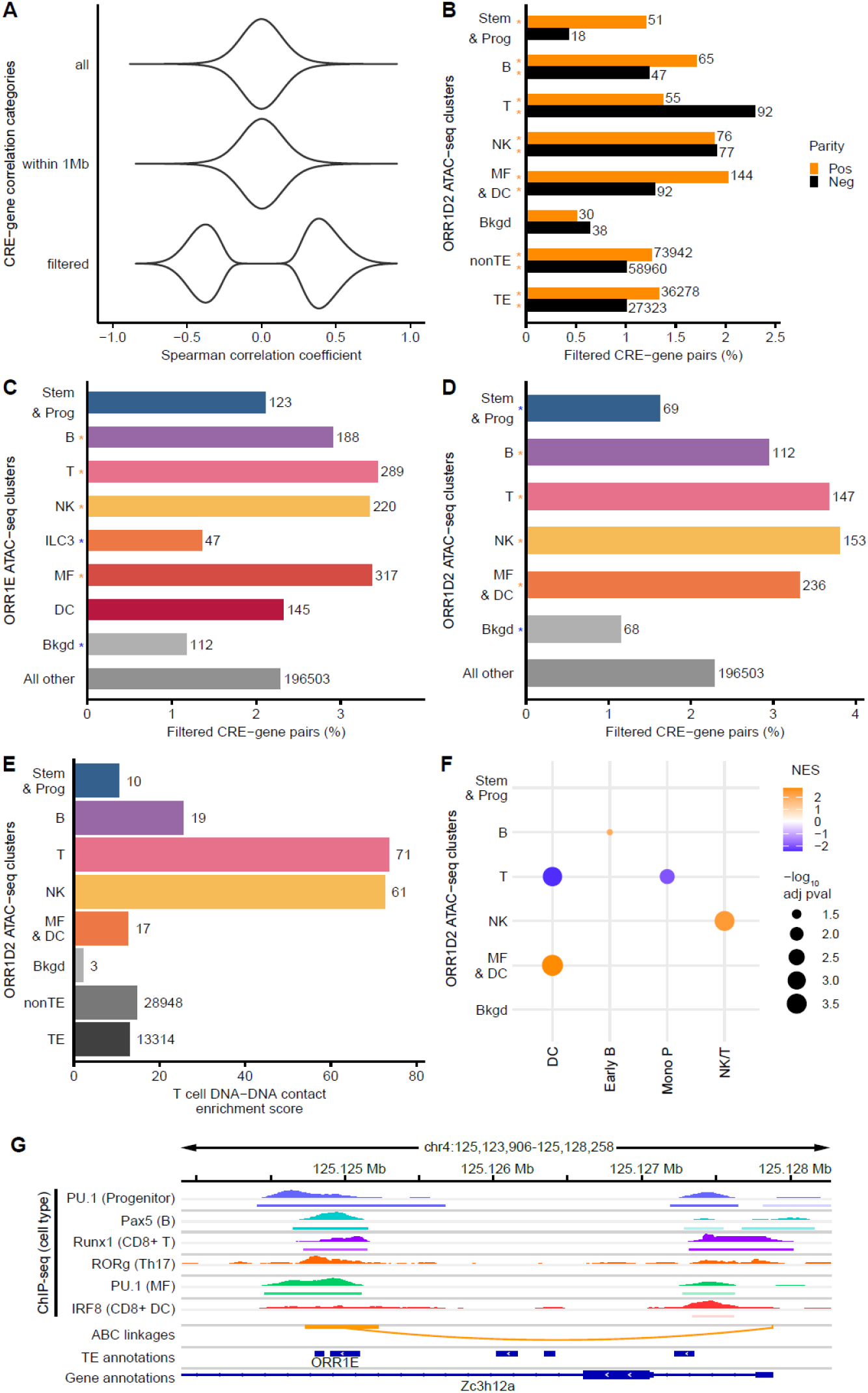
ORR1D2 are predicted to regulate hundreds of gene targets. (**A**) Violin plot showing distributions of Spearman correlation scores for peak-gene pairs for all pairs (∼462 million), pairs within 1Mb (∼8.7 million), or filtered pairs (198,729). (**B**) Barplot showing the proportion of filtered pairs for each ORR1D2 cluster and all other TE-derived or non-TE-derived peaks, split by positive (orange) and negative Spearman correlation coefficient (blue) with orange asterisk indicating significantly (Fisher’s Exact Test adjusted p-value<0.05) higher proportion of filtered pairs than the respective ORR1D2 background cluster.(**C – D**) Barplot comparing proportion of filtered pairs (both positively and negatively correlated) across ORR1E (C) and ORR1D2 (D) clusters against all other TE and non-TE pairs together. Significance (Fisher’s Exact Test adjusted p-value<0.05) is indicated with an orange asterisk for more filtered pairs than “All other”, or a blue asterisk for fewer filtered pairs than “All other”. (**E**) Barplot showing enrichment of DNA-DNA contacts in CD8+ T cells per ORR1D2 cluster and all non-TE and TE peaks. (**F**) Dotplot showing significant (adjusted p-value<0.05) GSEA results of ORR1D2 gene targets for cell type marker gene sets. (**G**) Browser track showing ChIP-seq binding data of PU.1 in progenitor cells, Pax5 in B cells, Runx1 in CD8+ T cells, RORg in Th17 cells, PU.1 in Macrophages, and IRF8 in CD8+ DCs, for an ORR1E locus directly contacting, as determined by the ABC model, the *Regase-1* (Zc3h12a) promoter in CD8+ T cells.

**Fig. S6:**
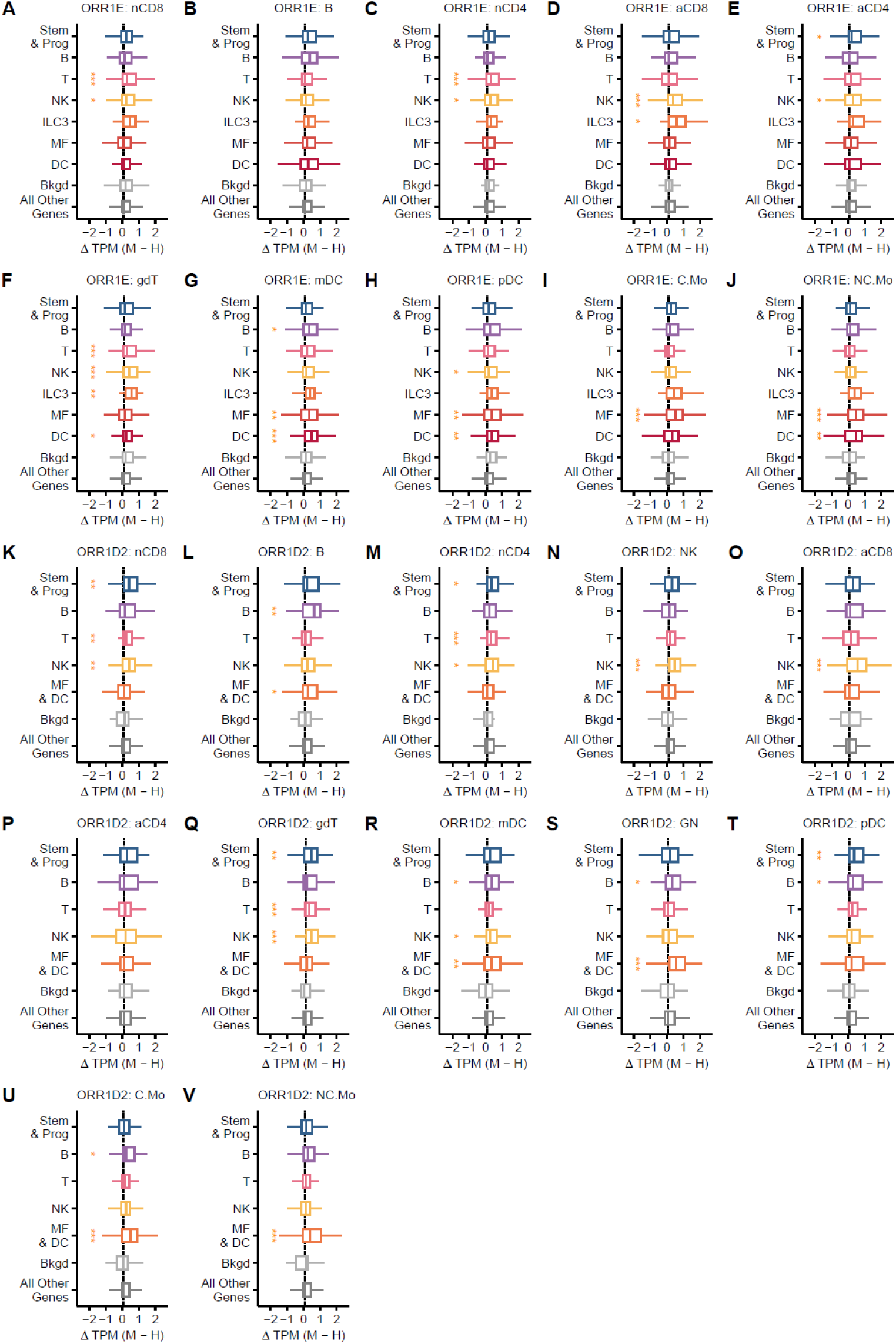
ODE-gene targets have cell-specific increases in gene expression in mouse compared to their human orthologs. (**A – V**) Boxplots showing distributions of changes in scaled gene expression (see Methods) for all positively correlated gene targets for ORR1E (A – J) and ORR1D2 (K – V) across all remaining orthologous cell types not present in Fig. 5A-B. nCD8 = naïve CD8+ T cells, nCD4 = naïve CD4+ T cells, aCD8 = activated CD8+ T cells, aCD4 = activated CD4+ T cells, gdT = gamma delta T cells, mDC = myeloid dendritic cells; pDC = plasmacytoid dendritic cells, C.Mo = Classical Monocytes, NC.Mo = Non-Classical Monocytes. Significance (Wilcoxon Rank Sum test compared to “All Other Genes”, adjusted p-value<0.05) is marked by asterisks (black = less; orange = greater), as follows: *p<0.05, **p<0.005, ***p<0.0005.

**Fig. S7:**
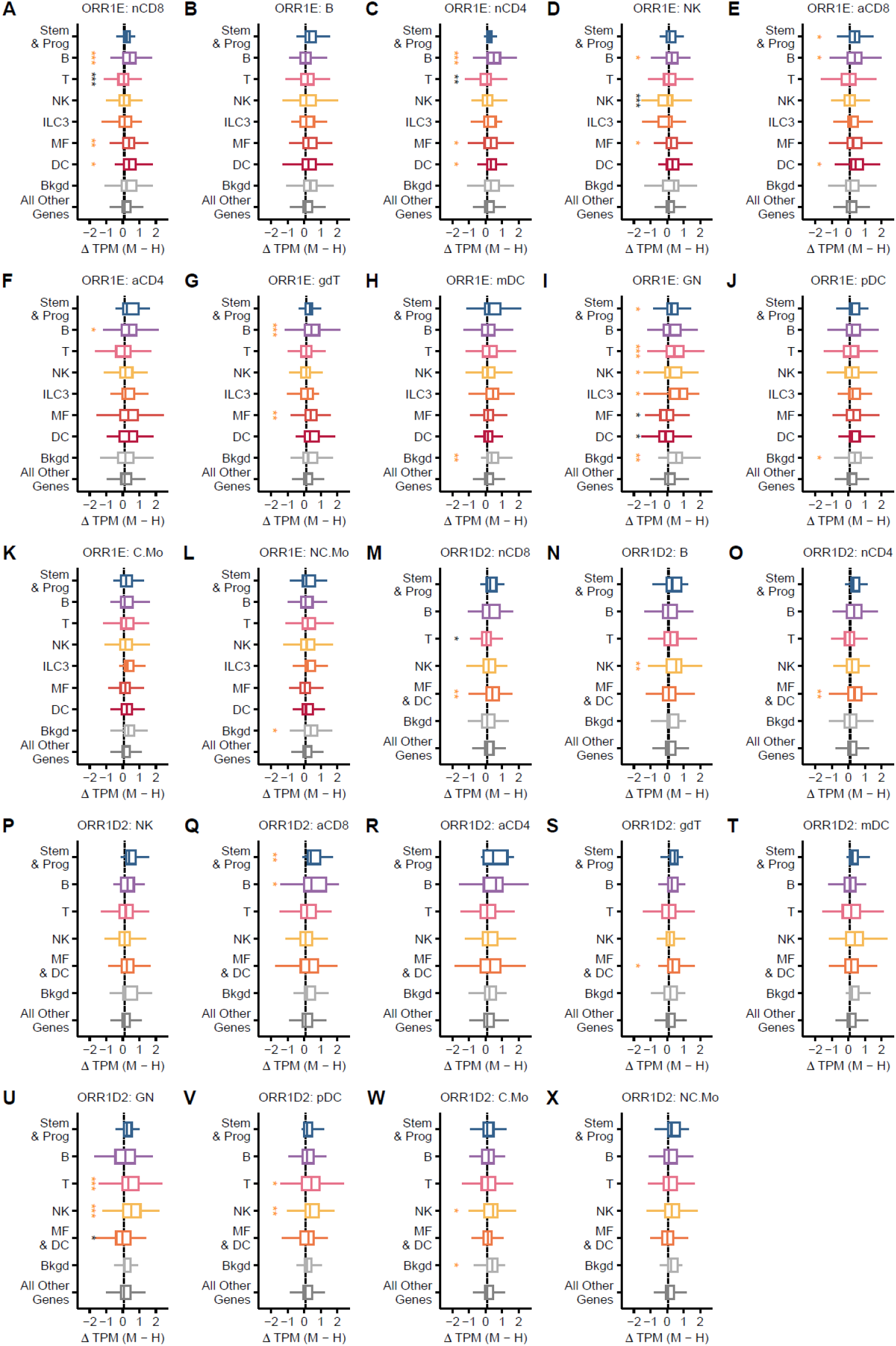
ODE-gene targets have cell-specific decreases in gene expression in mouse compared to their human orthologs. (**A – X**) Boxplots showing distributions of changes in scaled gene expression (see Methods) for all negatively correlated gene targets for ORR1E (A – L) and ORR1D2 (M – X) across all remaining orthologous cell types not present in Fig. 5A-B. nCD8 = naïve CD8+ T cells, nCD4 = naïve CD4+ T cells, aCD8 = activated CD8+ T cells, aCD4 = activated CD4+ T cells, gdT = gamma delta T cells, mDC = myeloid dendritic cells; pDC = plasmacytoid dendritic cells, C.Mo = Classical Monocytes, NC.Mo = Non-Classical Monocytes. Significance (Wilcoxon Rank Sum test compared to “All Other Genes”, adjusted p-value<0.05) is marked by asterisks (black = less and orange = greater) and *p<0.05, **p<0.005, ***p<0.0005.

**Fig. S8:**
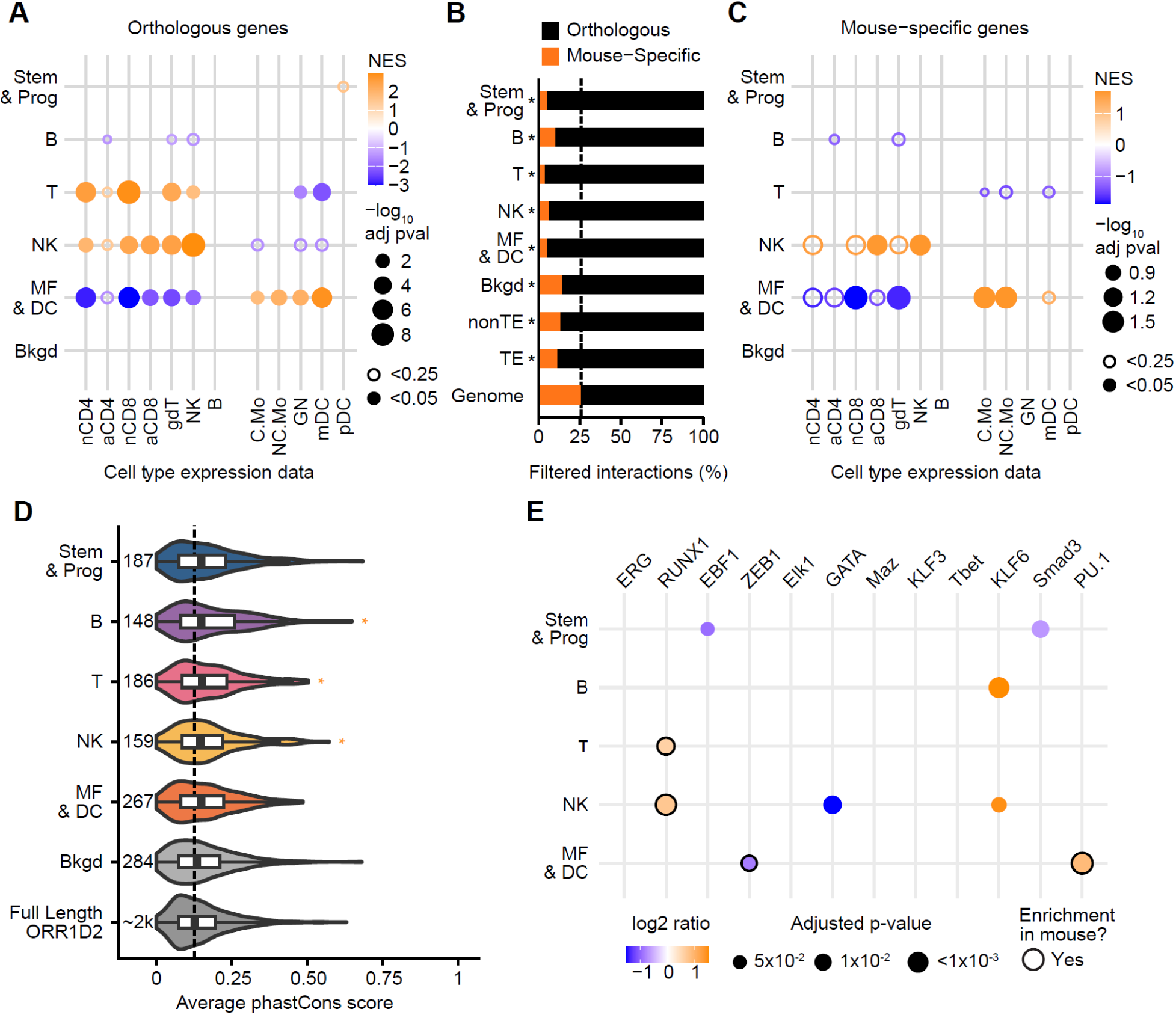
ORR1D2-gene pairs are associated with mouse cell type identity and have signatures of selection. (**A**) Dotplot showing significant (adjusted p-value<0.25 is open circle, adjusted p-value<0.05 is filled circle) GSEA results of ORR1D2 gene targets for gene sets that are more expressed in mouse compared to human, for orthologous (A) genes. (**B**) Barplot of proportions of peak-gene linkages for genes orthologous between mouse and human (black) or mouse-specific (orange). Asterisks indicate significantly (Fisher’s Exact Test adjusted p-value<0.05) lower proportions of mouse-specific target genes compared to genomic proportion. (**C**) As per panel (A) but using mouse-specific ORR1D2 gene targets. (**D**) Violin plot measuring significantly (Wilcoxon Rank Sum test, adjusted p-value<0.05) higher average phastCons scores for genes from ORR1D2 clusters (as per Fig. 2A) compared to full-length non-accessible ORR1D2s. Numbers indicate loci included per cluster. *p<0.05. (**E**) Dotplot of significant (Binomial test adjusted p-value<0.05) motif enrichment results from HOMER on orthologous ORR1D2 loci in rat across mouse clusters against orthologous background cluster in rat. Black outline indicates if cluster motif enrichment is also present in orthologous mouse elements.

**Fig. S9:**
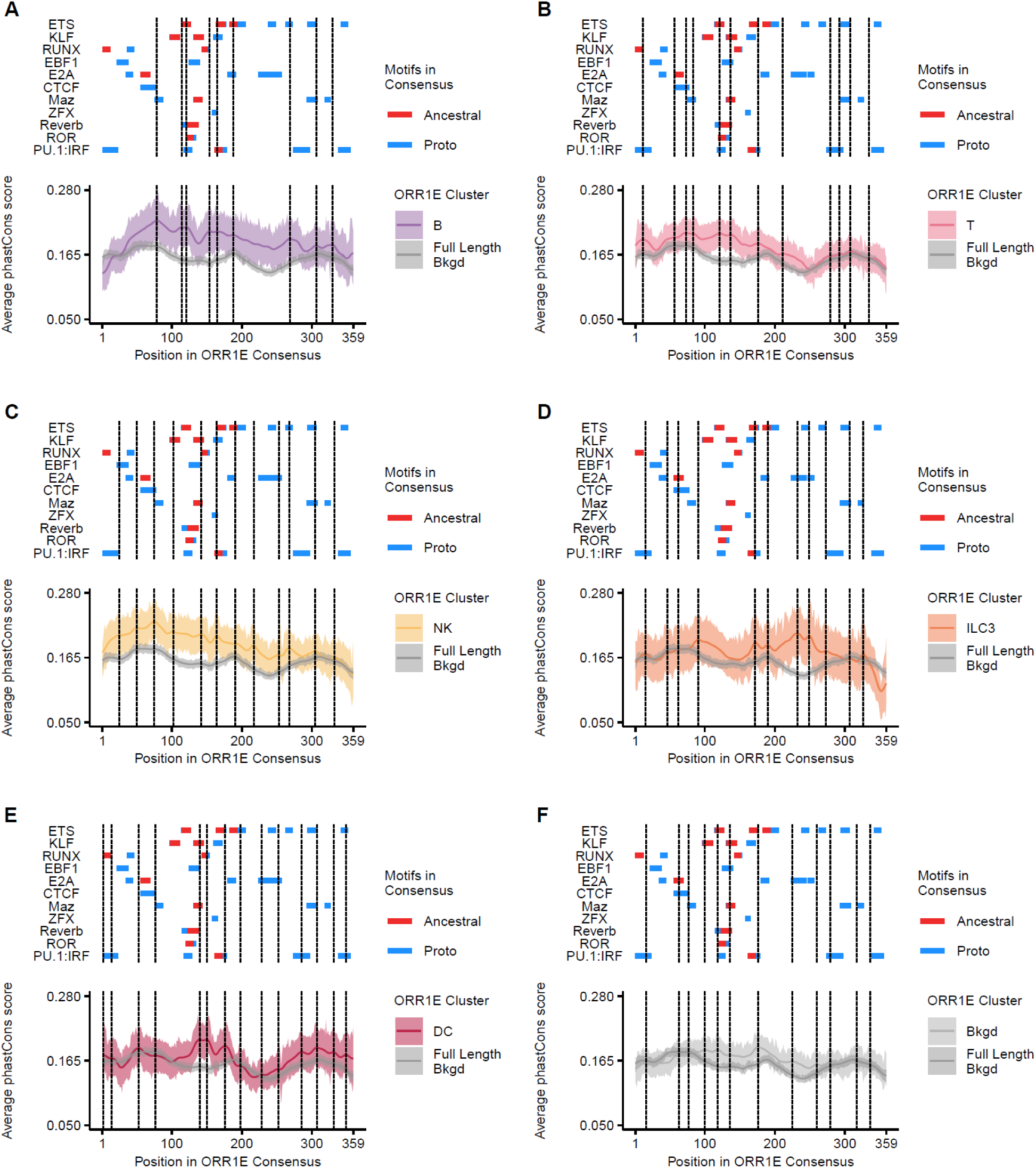
ORR1E conservation across clusters. (**A – F**) Visualization of matched motifs (Ancestral, red) or proto-motifs (Proto, blue) in ORR1E consensus for remaining enriched motif clusters from Fig. 2B not present in Fig. 6C-D. Density plots with 95% confidence interval of average phastCons scores for each ORR1E cluster and full-length background ORR1Es aligned to the consensus sequence. Boxed regions are representative local maxima of average phastCons scores.

**Fig. S10:**
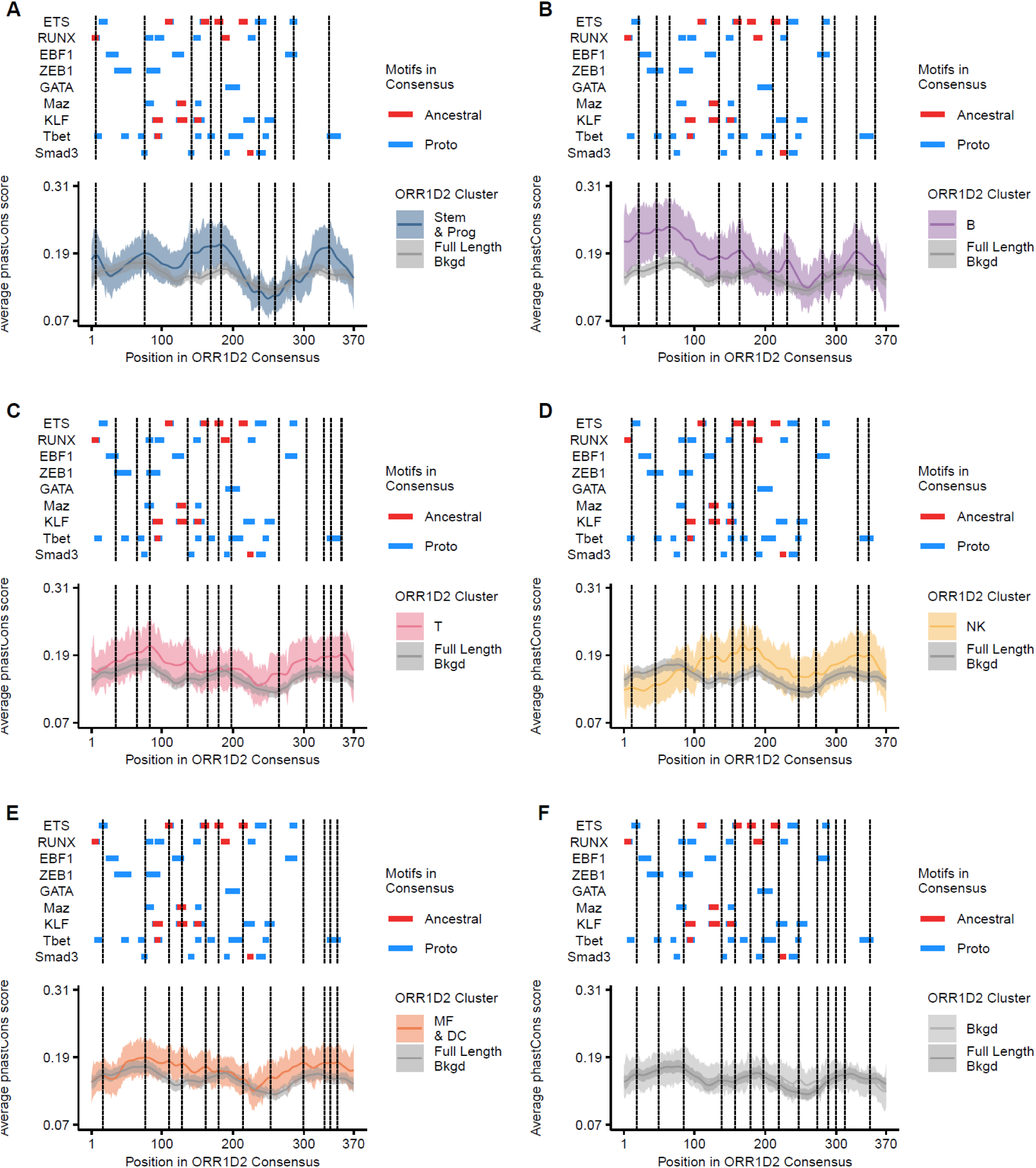
ORR1D2 conservation across clusters. (**A – F**) Visualization of matched motifs (Ancestral, red) or proto-motifs (Proto, blue) in ORR1D2 consensus for enriched motif clusters from Fig. S2A. Density plots with 95% confidence interval of average phastCons scores for each ORR1D2 cluster and full-length background ORR1D2s aligned to the consensus sequence. Boxed regions are representative local maxima of average phastCons scores.

