## Supplemental Data and Tables for "Pervasive *cis*-regulatory co-option of a transposable element family reinforces cell identity across the mouse immune system": tableS4_odeTopGeneContacts.txt

gene	cluster	factors_bound	relevant	function	source(s)
Zc3h12a	2		B/T/MF		Y		Is an RNase necessary for B cell response, activation of T cells and macrophages, and is involved in regulating many other interesting pathways	Bhat N et al 2021 (https://pubmed.ncbi.nlm.nih.gov/33822844/), Matsushita K et al 2009 (https://www.nature.com/articles/nature07924)
Cd86		2		B/T/MF		Y		ligand for CD28 expressed in B, DC, and monocytes (APCs) and is key for proper T cell activation via interacting with CD28	Esensten JH et al 2017 (https://pmc.ncbi.nlm.nih.gov/articles/PMC4932896/)
Bhlhe41	2		Prog/T		Y		controls development, BCR repertoire, and self-renewal of innate-like B-1a cells mostly via repression, other macrophage and B cell subsets, and is highly expressed in Th2 cells	Kreslavsky T et al 2017 (https://pmc.ncbi.nlm.nih.gov/articles/PMC5363839/#ref-list1), Bret C et al 2024 (https://link.springer.com/article/10.1007/s13402-024-00973-3)
Cd226		3		None		Y		Preferentially expressed in NK/T where it functions during T cell development and is critical for NK activation and killing by facilitating stable interactions with target cells	Liu K et al 2025 (https://www.mdpi.com/2218-273X/15/7/1007)
Nsg2		3		B		N		Only correlative association with immune cell infiltration and mostly serves as an AMPAR-binding protein that is required for normal synapse formation and/or maintenance 	Li X et al 2024 (https://pmc.ncbi.nlm.nih.gov/articles/PMC11527618/), Chander P et al 2019 (https://pmc.ncbi.nlm.nih.gov/articles/PMC6345199/)
Bicdl1	3		None		N		High expression is correlated with decreased patient survival and weakly associated with various immune cell infiltrations in human cancer	 Luo H et al 2023 (https://pmc.ncbi.nlm.nih.gov/articles/PMC10750784/)
Atf3		3		None		Y		Represses immune response genes post-activation in MFs, NKs, Th2s and Neutrophils	Ku H-C and Chen C-F, 2020 (https://www.frontiersin.org/journals/endocrinology/articles/10.3389/fendo.2020.00556/full)
Plbd1		3		DC		N		Only correlative association with human cancers but seems to play a crucial role in glioma cell proliferation, invasion and immunotherapy	Wei M et al 2024 (https://pmc.ncbi.nlm.nih.gov/articles/PMC11190780/#abstract1)
Prkcq		4		None		Y		Critical for T cell activation and is a well-studied gene	sources
Arap2		4		Prog/B		Y		Seems to be required for proximal signaling post TCR-stimulation and its loss is associated with reduced severity of autoimmune disease	Kim J-H, et al 2025 (https://www.frontiersin.org/journals/immunology/articles/10.3389/fimmu.2025.1556616/full#B22)
Tnk2		4		None		Y		Serves as a protein kinase upstream of CSK-binding protein (CBP) / PAG and helps restrict T cell activation. However, its role in other immune cells is unclear	Sridaran D and Mahajan NP 2025 (https://www.sciencedirect.com/science/article/pii/S0165614724002475#s0040)
Itpr2		4		Prog/B/T/MF	Y		KO is associated with decreased presence of memory T cells and increased lifespan	Ziegler DV et al 2021 (https://www.nature.com/articles/s41467-021-20993-z)
Fasl		4		None		Y		Is associated with both apoptosis and survival as a ligand to Fas	Yamada A et al 2017 (https://pmc.ncbi.nlm.nih.gov/articles/PMC5380675/#S3)
Bcl2a1b	4		None		Y		A member of the anti-apoptotic gene Bcl family and has been shown to be expressed in T cell development. Also generally other immune cells?	Schmitz I et al 2003 (https://academic.oup.com/intimm/article/15/10/1237/665986)
Samd3		4		T		Y		Is expressed in NK, memory CD8 and precursor exhausted T cells	Peters AE et al 2021 (https://onlinelibrary.wiley.com/doi/10.1002/eji.202149469)
Wipi1		4		None		N		Is involved in autophagy and other WIPI proteins are associated with human diseases	Almannai M et al 2022 (https://pmc.ncbi.nlm.nih.gov/articles/PMC9500159/)
Wfs1		4		Prog/B/MF	N		Mutations caused in this gene are associated with Wolfram Syndrome and has minimal evidence for immune-related function	Ding Y et al 2023 (https://www.frontiersin.org/journals/endocrinology/articles/10.3389/fendo.2023.1066320/full)
Tgfbi		6		MF/DC		Y		Disruption of TGFBI inhibited MF polarization to M2 phenotype but improved anti-tumor immunity	Zhou J et al 2023 (https://www.sciencedirect.com/science/article/pii/S0304383523004081)
H2-DMb1	7		Prog/B/T/MF/DC	Y		A gene part of the major histocompatibility complex preferentially expressed on MF and DC 
Tgtp1		7		Prog/B/MF/DC	N		Identified as an immune-related GTPase in neurons in response to positive-stranded RNA viruses, but not well studied elsewhere	Cho H et al 2013 (https://www.nature.com/articles/nm.3108)
Klrb1c	7		B/T		Y		Component of NK activation and NK killing and is also important in some T cell subsets
Klra1		7		B/T		Y		Component of NK activation and NK killing and is also important in some T cell subsets
Traf1		7		Prog/B/T/MF/DC	Y		Largely expressed in activated myeloid and lymphoid cells. Involved in activated/memory T cell survival	Edilova MI et al 2018 (https://pmc.ncbi.nlm.nih.gov/articles/PMC6305416/)
Gsap		3		Prog/B/MF/DC	N		Mainly assoicated with Alzheimer's via Amelyoid Beta production, but is suggested to play a role in innate immune functionality in rat KO experiments. Not enough to be relevant	Gwin MS et al 2023 (https://pmc.ncbi.nlm.nih.gov/articles/PMC10396227/)
Lpin1		4		None		Y		Is required for full phosphorylation of MAPKS ERK, JNK and p38 and thus c-Jun. Much known is focused on macrophages and is understudied in T and NK cells, but connection with c-Jun is notable	Balboa MA et al 2019 (https://www.sciencedirect.com/science/article/pii/S1388198119300952) 
F2rl1		4		T		Y		Plays an important anti-apoptotic role in T cell development as gene that encodes PAR2	Francis N et al 2017 (https://www.sciencedirect.com/science/article/pii/S1357272517302480)
Cd38		4		None		Y		Is a well-studied multifunctional transmembrane protein widely expressed in immune cells and is used as a cell activation marker for B cells	Glaria E and Valledor AF 2020 (https://pmc.ncbi.nlm.nih.gov/articles/PMC7017097/#sec7-cells-09-00228)
